# p125A (Sec23ip) couples COPII coat assembly with donor-acceptor membrane organization to facilitate tunnel-based traffic

**DOI:** 10.1101/2025.05.07.652703

**Authors:** Kimberly R. Long, Gazal Singh, Marie Villemeur, Nathalie Brouwers, Vivek Malhotra, Ishier Raote, Meir Aridor

## Abstract

COPII-coat proteins play a crucial role in generating small vesicles at endoplasmic reticulum (ER) exit sites (ERES). However, they also assemble at the necks of tunnels and tubules connecting the ER exit site (donor) to the ERGIC/cis-Golgi cisterna (acceptor). How can COPII support these two traffic mechanisms? Through cell-free reconstitutions, we have found that the apposition of donor and acceptor membranes is important for the assembly of the outer layer of the COPII coat (Sec13/31) but had minimal impact on Sar1-induced COPII inner layer (Sec23/24) recruitment. The expression of the adaptor protein p125A, which binds to phosphatidylinositol 4-phosphate (PI4P), Sec31 and Sec23, stabilized the contact between the donor and acceptor membranes and promoted COPII outer layer assembly. A p125A-chimera expressing a Golgi-targeted PI4P-binding domain could also support outer layer assembly. Notably, the C-terminal helical domain of Sec31A, which interacts with p125A, was essential for its assembly at ERES. In cells lacking p125A, the assembly of the COPII outer layer was selectively destabilized. Transcriptome and secretome analyses reveal selective adjustments to extracellular matrix remodeling and collagen secretion, which corresponded with selective inhibition of fibrillar collagen traffic from the ER. Thus, p125A connects the outer COPII layer at ERES with PI4P-rich ERGIC/cis-Golgi membranes, coordinating COPII assembly with tunnel-driven collagen traffic.

## Introduction

The activation and recruitment of the small GTP binding protein Sar1 to endoplasmic reticulum exit sites (ERES) initiates a sequential assembly process, where the COPII coat inner layer Sec23/Sec24 is recruited followed by the assembly of outer layer subunits Sec13/Sec31 on a Sar1-Sec23 platform (1). The linking between the two coat layers stimulates GTP hydrolysis allowing for uncoating while facilitating membrane fission(2, 3) (4–6). This assembly process generates small vesicles in vitro from model membranes(7, 8). While small COPII vesicles have been observed in yeast, their visualization in metazoan cells is limited. Additionally, these vesicles are unable to export bulky molecules including extracellular matrix (ECM) proteins, a crucial requirement for animal multicellularity. Recent molecular analyses, combined with nano-resolution microscopy, support the involvement of tunnels or tubules in cargo export from the ER (9, 10). There is evidence suggesting that tunnels form between the donor ERES and the acceptor membranes (ERGIC/cis Golgi) through interactions of the TANGO1 family of ERES proteins with tether and fusion factors (11, 12). In fact, yeast, which also utilizes a similar contact-based traffic mechanism, suggests a simpler form where ER COPII buds transiently connect with Golgi membranes to mediate traffic in a “hug and kiss” type reaction (13).

COPII proteins play a dual role in cargo export at the ER: they can coat small vesicles entirely and also participate in the export of the bulky ECM through tunnels (2). Studies have revealed COPII coats at the necks of tubules emerging from ERES (9, 10). The question arises: how can the vesicle forming COPII coat support tunnel-based traffic? Inter-compartment tunnels could form if a budding carrier fuses to the next compartment before the completion of fission.

However, if and how COPII assembly is coordinated with the recruitment of acceptor membranes remains an open question. p125A (Sec23ip) could play a role in this step of tunnel biogenesis. p125A is a Sec31-binding adaptor and a stoichiometric component of the Sec13/31 complex in the cytosol (14, 15). It interacts with Sec23 upon membrane recruitment, thus physically linking the inner and outer coat layers. This interaction facilitates the coordination of COPII assembly at the ER exit sites (ERES). Moreover, p125A binds PI4P and supports Sec13/31 assembly at ERES (14, 16). Since PI4P is enriched on acceptor ERGIC and cis Golgi membranes, p125A may be in a unique position, simultaneously interacting with COPII at the ERES and PI4P at downstream compartments. We posit that these activities may coordinate between tunnels/tubules formation and coat-mediated cargo transport. PI4P regulates ERES assembly, adjusts ERES to acute cargo load, and regulates general cargo traffic from the ER (17–19). The utilization of PI4P in ER to Golgi traffic is evolutionarily conserved, regulating COPII vesicle fusion in yeast, which employs transient membrane contacts to support a “hug and kiss” based traffic mechanism (20) (13).

We investigated how COPII assembles at connected ERES-ERGIC-cis Golgi secretion hubs. Our findings demonstrate that p125A plays a role in coupling donor-acceptor membrane contacts with coat assembly, by simultaneously binding Sec31 at the ERES and ERGIC/Golgi membranes. The interaction with p125A facilitates coat outer layer assembly through PI4P binding. Deletion of p125A or removal of its binding site on Sec31 leads to the selective destabilization of COPII outer layer assembly at ERES, resulting in the selective inhibition of tunnel-based traffic from the ER. We propose that p125A serves as a link, coupling acceptor membrane contacts with outer layer coat assembly and GTPase activation, thereby coordinating coat functions with tunnel-based traffic.

## Results

### The recruitment of COPII inner and outer layers can be uncoupled

Multiple redundant mechanisms and proteins maintain ERES-ERGIC function and organization. To overcome this complexity, we designed a ‘rundown-type reaction’, where cells are permeabilized and then incubated for varying lengths of time before assaying for ERES-ERGIC machinery and function (21). Following permeabilization, cytosolic factors will be gradually released from membranes, diluted, and removed from the cells, leading to a rundown outcome, where dynamic and active processes are reduced progressively. At later times, only the most robust and stable activities will remain, allowing for the dissection of key traffic intermediates. A similar approach was instrumental in analyzing regulated secretion in mast cells (21).

We adopted a rundown protocol in which permeabilized cells HeLa were pre-incubated at 32°C in the absence of cytosol, followed by the addition of rat liver cytosol with recombinant Sar1-GTP (Sar1^H79G^). The cytosol would provide COPII and regulatory proteins required to reconstitute COPII assembly. Under control conditions, Sar1-GTP induced recruitment of both COPII layers to defined ERES, as expected (Fig. 1). Under the rundown conditions, the recruitment of the coat inner layer (Sec24C) was still effective (Fig. 1B and C-D) but interestingly, the recruitment of the COPII outer layer Sec31 (Fig. 1A, C, H and Fig. 4E) was quantitatively and robustly inhibited whereas the recruitment of Sec24C was largely unaffected (Fig. 1C). We have monitored Sec31 recruitment using two antibodies, a rabbit one that recognizes both human and rat Sec31A (Fig. 1A, 1C, 1H, Fig. 2 and Fig. 4E) and a monoclonal antibody that preferentially recognizes human Sec31A (Fig. 1D) (22).Thus, COPII outer layer proteins are recruited from cytosol but also retained at ERES through cell permeabilization and washes and stabilized by Sar1 activation. In a similar manner the recruitment of Sec13 was inhibited in rundown conditions (Fig. 1E, Fig. S1A). Under the permeabilization conditions, microtubule network disruption leads to changes to the ER network(23). However, ER morphology was similar in control and rundown conditions as marked by an ER localized reporter, a Tac chimera that presents the cytosolic C-terminus of gp25L containing a di-lysine retrieval / retention motif, (Tac-gp25l, Fig. 1E). COPII was recruited by Sar1-GTP from cytosol to ERES showing strong colocalization of Sec31A with Sec16A, that was perturbed when monitored in cells depleted of Sec16A prior to permeabilzation and reconstitution of coat assembly (Figs. 1F and S1B). ERES were not disrupted in rundown conditions as monitored by Sec16A localization (Figs 1F and S1B). We further expressed GFP tagged Sec16A to monitor the recruitment of the coat inner layer at ERES. In these cells, recruited Sec24C juxtaposed and overlapped with EGFP-Sec16A at ERES both in control and rundown conditions (Fig. 1G). Overall, the results suggest that under rundown conditions, the COPII inner layer is effectively assembled at ERES while outer layer assembly is perturbed (Fig. 1A-E and G-H and Fig. S1A). We confirmed that Sar1 activation was unaffected under rundown conditions by showing that Sar1-dependent ERES membrane tubulation was unaffected in incubations carried out only in the presence of Sar1-GTP [(3, 24), not shown]. Since the recruitment of the COPII outer layer at ERES was selectively inhibited, it appears that rundown incubations uncouple the two coat layers.

**Figure 1.**
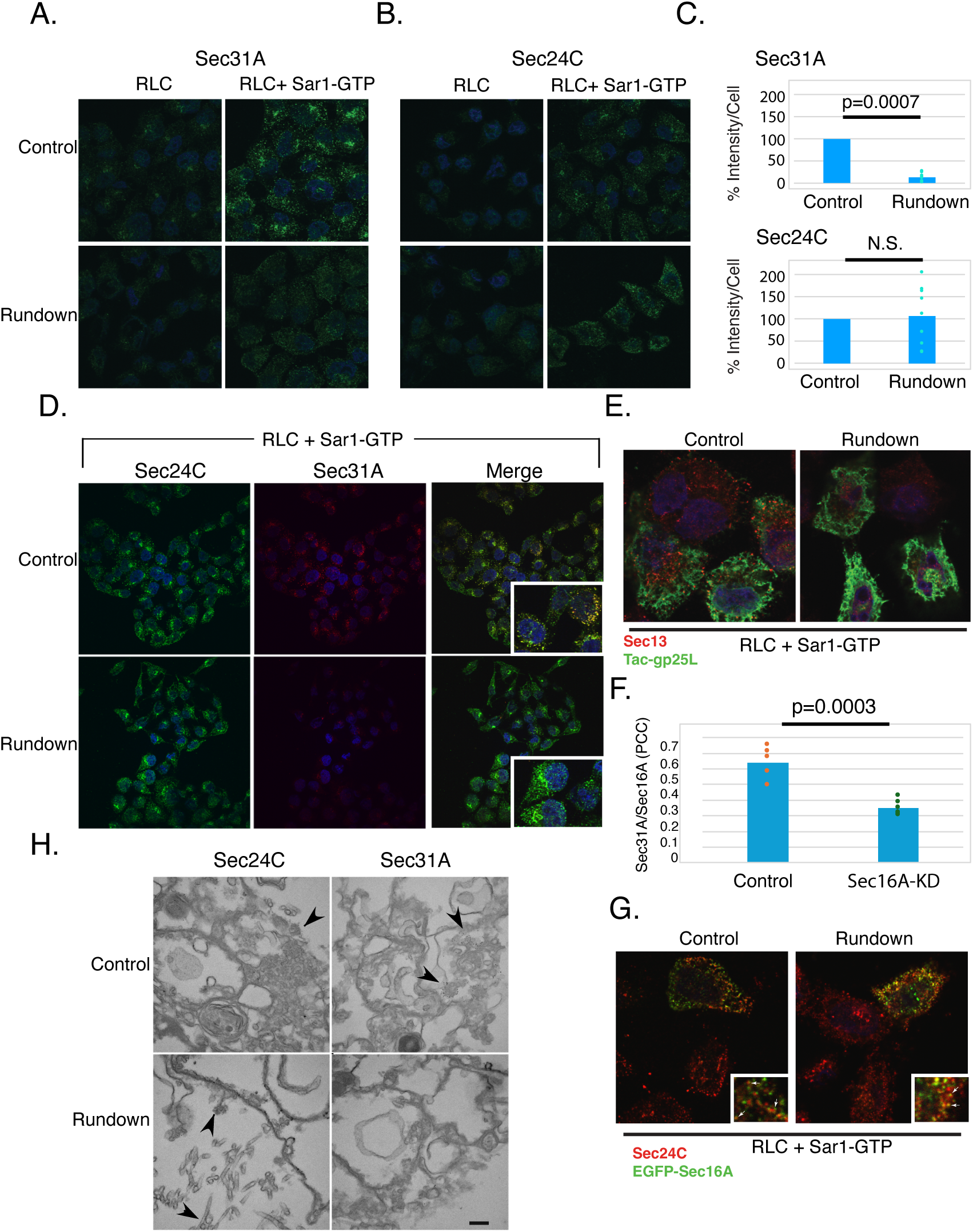
The assembly of COPII outer layer at ERES is inhibited after a “rundown incubation. **A-B.** Permeabilized HeLa cells were either pre-incubated in a reaction buffer lacking nucleotides, cytosol or SAR1 proteins for 15 min at 32°C (rundown) or not (control). Subsequently, a full transport cocktail containing rat liver cytosol (RLC) and SAR1-GTP (hamster SAR1a-H79G, 5μg/220μL) were added as indicated. The reaction was carried out for 15 minutes at 32^°^C, the cells were washed, fixed and analyzed for Sec31(using a polyclonal anti Sec31A antibody, **A**.) or Sec24C (**B.**) at ERES as we described (3) **C.** The intensity of Sec31A or Sec24C was quantified from 9 fields of cells (20-60 cells per field, each field represented by a circle) derived from 3 to 4 independent experiments. Controls were defined as 100 percent and results analyzed using two-tailed t-test. **D.** Control and rundown reactions were as described in A in the presence of SAR1-GTP and RLC. Fixed cells were processed for IF using anti Sec24C and Sec31A (monoclonal) antibodies as indicated. **E.** Permeabilized cells expressing Tac-gp25l (ER marker) were incubated under control or rundown conditions (as in A) with SAR1-GTP and RLC, fixed and stained for Sec13 (red) and Tac (green). **F.** Analysis of Sec31A recruitment to ERES (marked with anti Sec16A antibody) in permeabilized control or Sec16A-KD cells (staining control) supplemented with Sar1-GTP and RLC as in A was analyzed from 6 fields of cells per condition (circles) from two independent experiments using Pearson Correlation Coefficient (PCC) (shown with two tailed t-test) **G.** Cells expressing GFP-tagged Sec16A were permeabilized and incubated as in A with RLC and Sar1-GTP, fixed and stained for Sec24C (red). Arrows in inserts indicate juxtapose labeling of recruited Sec24C with ERES markers in control and rundown conditions**. H.** Reconstitution of COPII recruitment was conducted as described in A. The cells were fixed and labeled with antibodies to Sec31 (polyclonal) and Sec24C, further fixed and processed for EM (3, 26). Bars are 5μm in A and 200nm in E. Arrows in E point to gold labeling of Sec24C or Sec31 at ERES as indicated.

**Figure 2.**
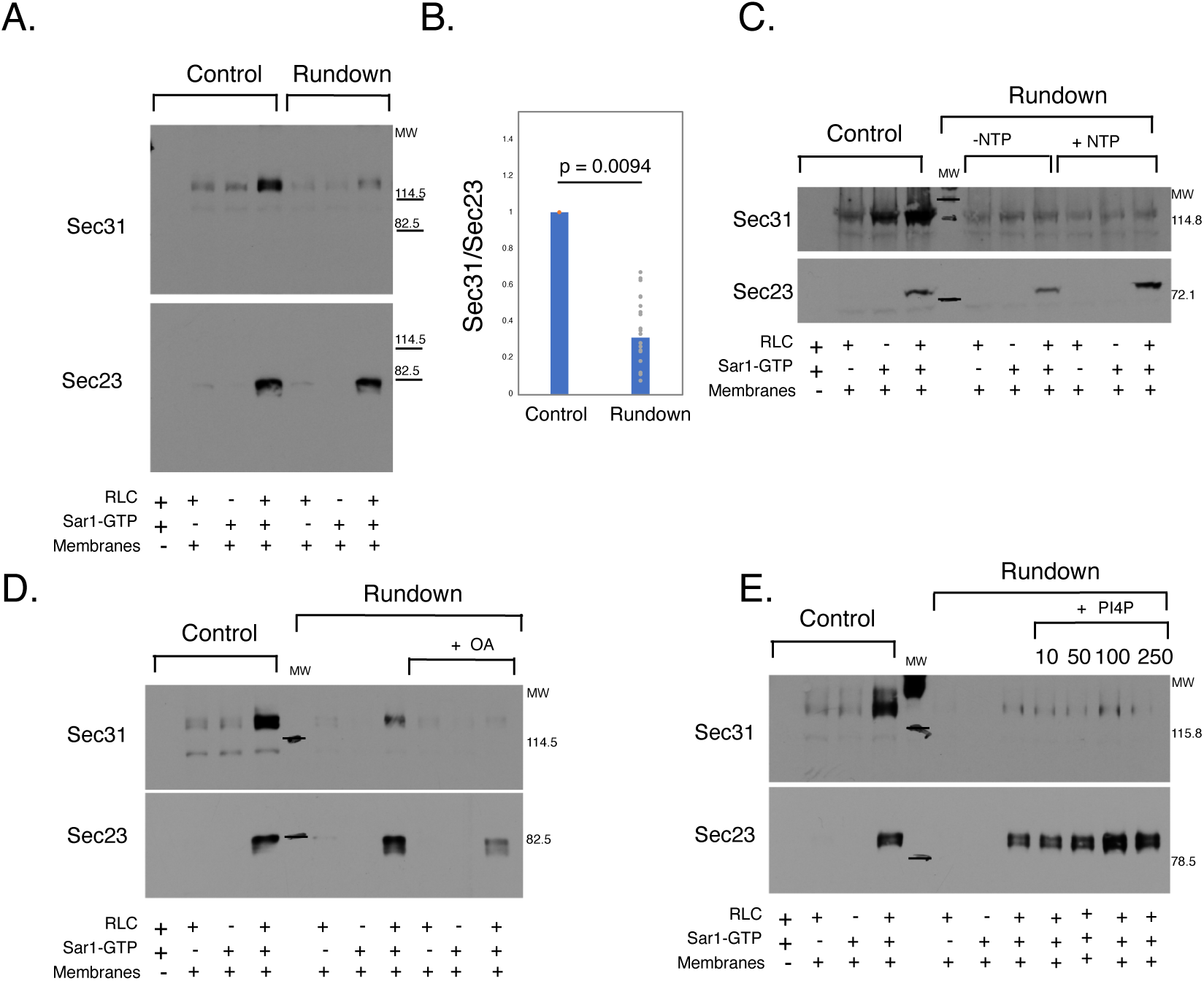
The recruitment of COPII outer layer to ER membranes is inhibited after a rundown incubation. **A.** Control HeLa microsomes or ones subjected to a pre-incubation rundown protocol as in Fig. 1 were subsequently incubated with SAR1-GTP (300ng/60μL) and/or RLC as indicated. The microsomes were salt-washed and analyzed for coat recruitment using SDS-PAGE and western blots that were probed, stripped and re-blotted using antibodies against Sec31 and Sec23 as indicated (3, 17, 25).**B.** The density ratios between recruited Sec31 and Sec23 per lane were quantified for control and rundown incubations carried with Sar1-GTP and RLC, using 22 independent experiments. For each experiment control ratio was set at 100% and experiments were than averaged. The outcome of individual experiments is presented with the overall average, (two-tailed t-test between groups is shown). **C.** Microsomes were subjected to rundown conditions, where the rundown incubation was done with or without added NTPs (ATP regenerating system and GTP) as indicated. Rundown or control microsomes were subsequently incubated with nucleotides in the presence of RLC and SAR1-GTP as indicated and analyzed for Sec31 and Sec23 recruitment. **D.** Microsomes were subjected to rundown conditions in the presence or absence of Okadaic Acid (OA, 2.5μM) as indicated. Subsequently, control or rundown microsomes were incubated in the presence of RLC and SAR1-GTP (as in A) and the recruitment of Sec23 and Sec31 was determined. **E.** Microsomes were subjected to rundown conditions in the presence or absence of increasing concentrations of PI4P micelles (in μM) as indicated (17). Subsequently, control or rundown microsomes were incubated with SAR1-GTP and RLC as indicated and the recruitment of Sec23 and Sec31 was determined by western blot with specific antibodies as in A.

### Sec13/31 recruitment to microsomes is inhibited in rundown conditions

The morphology-based coat assembly assays suggest that the coat inner layer was effectively assembled at ERES under rundown conditions while the outer later was not – thus Sec23/24 concentration at ERES was independent of Sec13/31 polymerization (Fig. 1). We needed an independent robust quantitative assay to further evaluate large populations of ERES and determine the magnitude of the observed rundown effect in a manner that goes beyond morphological analysis. We therefore utilized isolated salt-washed microsomes to reconstitute membrane binding of COPII proteins following Sar1-activation (25, 26). Incubation of microsomes with cytosol and Sar1-GTP led to the recruitment of both inner and outer COPII layers as monitored by concomitantly following Sec23 and Sec31 on the same western blots (Fig. 2, 2A). The reaction was dependent on the addition of cytosol with Sar1-GTP and membranes as previously described (3, 17, 25). These assays started with salt washes to extract peripherally attached proteins from microsomes. Therefore, we could selectively focus on recruitment of cytosolic Sec31A (Fig. 2A). In agreement with our quantitative microscopy-based analysis in permeabilized cells (Fig. 1C), subjecting microsomes to the rundown protocol led to selective inhibition of outer layer recruitment whereas the recruitment of the inner layer was only slightly affected (Fig. 2A-E). The microsome-based assay, which averages millions of cells and ERES, was further used to quantify the overall rundown effect on COPII outer layer recruitment (Figs 1 and 2). For the analysis, we now quantified the ratio between inner and outer layer recruitment on membranes in control and rundown conditions using 22 independent experiments. For each experiment, we defined the ratio between inner (Sec23) and outer layer (re probed for Sec31) per sample/lane on the same western blot, with Sec23 providing an internal control. The results showed an averaged inhibition of 70% in the recruitment of the coat outer layer when compared with inner layer assembly (Fig. 2B) in agreement with morphological analysis (Fig. 1C). Collectively, the results suggest that the rundown incubation led to quantitative uncoupling between COPII inner and outer layers at ERES.

### The rundown response is ATP-independent

We considered other ERES components that might be affected by the rundown and have a role in this coupling of inner and outer layers. Since the experiment used fresh cytosol and rundown cell membranes, the component cannot be soluble cytosolic proteins, but it could be membrane-associated peripheral protein. Sec16A is a peripheral membrane protein at ERES that partitions between cytosol and membranes (27, 28) and is known to regulate outer layer membrane recruitment, competing with Sec31 for Sec23 binding and inhibiting GTPase activation(29, 30). To test its role in the rundown assay, we depleted Sec16A using siRNA and assayed whether this rescued Sec13/31 recruitment after rundown. However, we observed no rescue – Sec16A depletion did not relieve the inhibition of Sec13/31 assembly after the rundown (Fig. S1B).

ATP and kinase activities regulate COPII assembly (31) and might contribute to the reduced recruitment of the coat outer layer during the rundown assay. To examine if kinase activities are modified during rundown incubations, ATP regeneration system and GTP (NTPs) were added or omitted during rundown incubations of washed microsome membranes. Subsequently both NTPs were re-added along with cytosol and Sar1-GTP to support the coat assembly reaction and the recruitment of both coat layers was monitored. While Sec23 recruitment was slightly enhanced by the presence of NTPs during the initial rundown incubation, Sec31 binding remained inhibited, suggesting that the phosphorylation status of membrane components is not the cause for the selective inhibition of Sec13/31 recruitment (Fig. 2C).

Another possible option was Phosphatase 2A (PP2A), as it has been shown to function in an early step in ER-exit (32). Treatment of membranes during rundown incubations with the PP2A inhibitor okadaic acid (OA), inhibited the recruitment of the coat inner layer (Sec23) and did not reverse the inhibition of outer layer assembly (Fig. 2D).

We next tested whether the phospholipid PI4P, which supports COPII assembly on liposomes and regulates ERES assembly in cells, might rescue the rundown effect (14, 17, 18). Addition of short chain PI4P micelles to microsomes during rundown incubations increased the recruitment of Sec23 to membranes in a dose dependent manner (Fig. 2E)(8). However, while these conditions also slightly elevated the recruitment of Sec13/31, the selective inhibition of Sec13/31 assembly in rundown conditions was not reversed (Fig. 2E).

### The helical C-terminal domain of Sec31 is utilized for assembly at ERES

Having ruled out these additional components or conditions, we considered the possibility that the binding itself between the two coat layers is regulated directly. The links between coat layers are established by the C-terminus of Sec31A, which contains a disordered proline-rich linker and a C-terminal α-solenoid-like helical domain. Within the proline-rich domain is a short segment containing the GAP-active tryptophan and asparagine residues (WN), which binds to an extended interface generated by Sec23 and Sar1 (6). These interactions are strengthened by the binding of a triple proline motif (PPP) to the gelsolin domain of Sec23 (33)

We analyzed the role of the helical C-terminus in supporting assembly at ERES. We truncated Sec31A to remove the C-terminal helical domain, or the domain along with the preceding proline-rich segment that links to the GAP-active Sar1-Sec23 binding WN domain (Fig. 3B) and verified that the WN-segment binds Sec23 in GST pull down assays (not shown). Because the analyzed constructs preserve the architectural core of Sec31A and can polymerize with ERES-localized endogenous Sec31A, we utilized 3’-UTR targeting siRNA to deplete endogenous Sec31A and replaced it with fluorescently tagged (FP-tagged) rSec31A (rat) proteins (Fig. 3A)(34). Both wildtype and truncated forms of FP-rSec31A were well expressed in control and depleted cells, suggesting that the truncations did not compromise overall protein stability (Fig. 3A and E). For morphological analysis, we permeabilized cells prior to fixation to wash-out overexpressed cytosolic Sec31A and focused on ERES-associated proteins. CFP-rSec31A localized to ERES in control cells, and in Sec31A-depleted cells (Fig 3C). In contrast, both truncated forms of Sec31A showed abnormal localization. In control cells, the FP-rSec31A truncations clustered in few discrete enlarged patches. In Sec31A-depleted cells, both truncated forms largely lost ERES localization and diffused staining was observed at different expression levels within analyzed fields, yet few cells still showed discrete large patch-like morphology (Fig. 3C). Therefore, the truncated Sec31A proteins retain the ability to interact with endogenous Sec31A, most likely through the polymerization core as seen in non-depleted cells, but the C-terminal helical domain of Sec31A was required for proper assembly at ERES. The results suggest that the Sec23-binding WN linker and the polymerization core of Sec31 are insufficient for proper nucleation of the coat outer layer at ERES, the C-terminal helical domain is also required.

**Figure 3.**
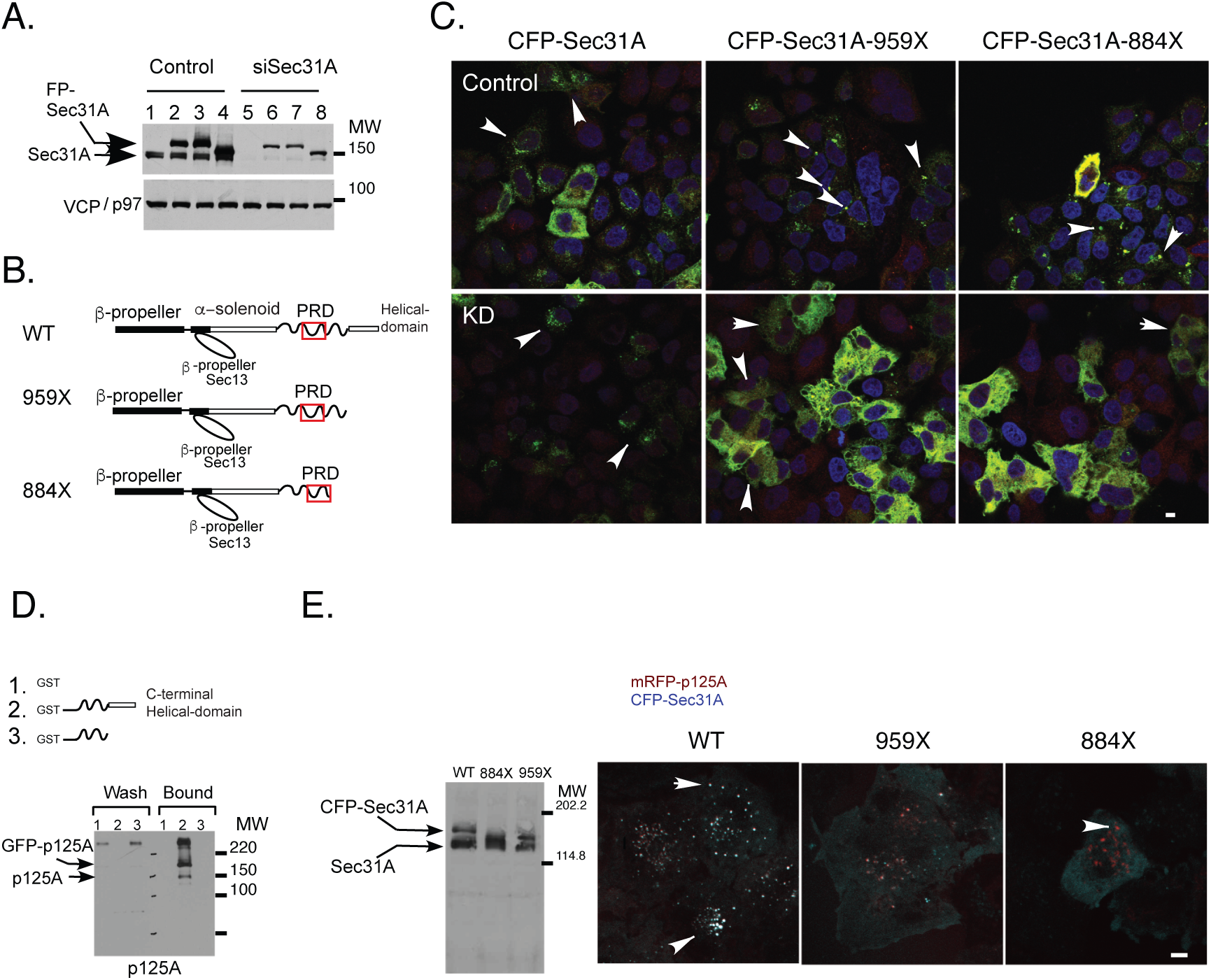
Sec31A requires the C-terminal α-Solenoid-like helical domain for effective assembly at ERES and for interactions with p125A. **A.** Control or Sec31A-depleted cells were transfected with resistant clones of YFP-rSec31A (lanes 2 and 6), CFP-rSec31A (3 and 7) or CFP-rSec31A-884X (4 and 8) and probed with Sec31A antibody to detect endogenous and tagged Sec31A. p97/VCP served as loading control. **B**. Schematic representation of rat Sec31A domains showing wild type (WT), a truncation of the C-terminal α-solenoid-like helical domain (959X rSec31A) and of the C-terminal helical domain and the proline-rich link to the Sec23-binding fragment (884X). The GAP active Sec23-binding fragment is labeled as a red box. **C.** Control or Sec31A depleted cells were transfected with CFP-rSec31A, CFP-rSec31A-959X or CFP-rSec31A-884X and the localization of Sec31A (red) and CFP (green) was defined by IF using specific antibodies. Arrows point to cells showing FP-Sec31A coated ERES or diffuse appearance in cells expressing the indicated FP-Sec31A truncations **D.** GST (1), GST-hSec31a-1041-1220 (2, proline-rich linker and terminal α-solenoid) or GST-hSec31a-1041-1113 (3, proline-rich linker), incubated with HeLa cell-lysates expressing GFP-p125A, collected on GS beads and probed with p125A antibodies. Note aggregation of p125A proteins under these conditions and the binding of both GFP tagged and endogenous p125A by the α-solenoid containing fragment. **E.** Left, the expression of CFP-rSec31A, or truncated forms (959X and 884X) as indicated, probed with polyclonal Sec31 antibody. Right, the cellular localization of co-expressed mRFP-p125A, (red) and CFP-rSec31A WT, 959X and 884X proteins (blue) as indicated (merged images are shown). Arrows point to Sec31-p125A colocalization at ERES, or lack of in cells expressing Sec31A truncations. Bar is 10μm.

### The C-terminus helical domain of Sec31A binds p125A

Yeast Sec31p utilizes multiple PPP motifs that compete for a single binding site on Sec23 or can also bind adjacent Sar1-Sec23/24 complexes to organize the inner layer in scaffolds (35, 36). These interactions are facilitated by the binding of the C-terminal helical domain onto the central α-solenoid dimerization arms of the Sec13/31 rhomboidal lattice stabilizing cage assembly. Inner-outer layer linking is further supported by multiple charge-based interactions. The reduced number of PPP motifs and reduced dependency of Sec31/13 rhomboidal assembly on the C-terminal α-solenoid in mammals suggest that metazoans and yeast differ in their mode of coat layer linking. Adaptors evolved in mammals to facilitate the linkage between the two coat layers (2, 35). Of these, p125A (Sec23ip) is a stoichiometric component of the cytosolic Sec13/31 complex that binds Sec23 upon membrane recruitment. We performed pull-down assays using GST-tagged Sec31A domains that affected assembly at ERES and examined interactions with p125A. p125A selectively interacted with a fragment containing the C-terminal α-solenoid-like helical domain, but not with the remaining proline-rich linker (Fig. 3D). Importantly, under the analyzed conditions, the C-terminal helical domain-containing fragment depleted both endogenous and expressed GFP-tagged p125A from cell lysates (Fig. 3D). To verify that p125A interacts with the C-terminal helical domain of Sec31A, we also co-expressed mRFP-p125A together with WT CFP-rSec31A or the Sec31A truncated forms in HeLa cells. Over expression of p125A leads to clustering of cargo containing ERES that collect COPII as well as ERGIC membranes and can report on p125A-Sec31 interactions(14, 37). The overexpression of mRFP-p125A effectively collected CFP-rSec31A at enlarged ERES (Fig. 3E). In contrast, the truncated forms of Sec31A did not associate with p125A at ERES but remained cytosolic, confirming that the C-terminal helical domain of Sec31A binds p125A (Fig. 3E).

Having identified p125A as an adaptor that could have an important role in this process, we could now propose hypotheses for how COPII-layers are coupled and what changes during the rundown that leads to a loss of outer layer recruitment. In addition to the ability to bind sec31A, p125A is a PI4P-binding adaptor. Interestingly, PI4P is enriched at ERGIC and Golgi membranes. This led us to hypothesize that p125A may bind PI4P that is enriched on acceptor ERGIC / cis Golgi membranes and that these membranes are unavailable for p125A binding in rundown conditions due to a breakdown of ERES-hub like organization (14, 16).

### Spatial organization of ERES-hub membranes is disrupted in rundown incubations

ERES are assembled from COPII coated ER membrane buds that are connected to the ERGIC by tunnels. In human cells, these connections appear as single tunnels coated with a COPII collar linked to ERGIC tubules(9). It is possible that connectivity is stable. Alternatively, a dynamic process of budding, fission and fusion is synchronized to generate tunnels (Fig. 7A). In drosophila, ERES-ERGIC tunnels consist of 1-3 fused vesicles supporting transient connectivity(38). To facilitate tunnel-based traffic, delayed fission by coordinating COPII outer layer recruitment with membrane organization, along with increased tethering and fusion may be necessary.

We analyzed the localization of exogenously added Sec24C recruited by Sar1-GTP in relation to that of acceptor membranes (ERGIC53, ERGIC) or Giantin (cis Golgi) in control and rundown conditions. Recruited Sec24C partially overlapped both with ERGIC53 membranes at peripheral ERES and with Giantin at the juxtanuclear Golgi region (Fig. 4A and not shown). In contrast, this overlap was lost under rundown conditions. The overall ERGIC53 signal was reduced during the rundown incubation, attesting to a loss of ERGIC membranes from permeabilized cells, as previously described [(39) and see below]. The remaining ERGIC53 signal localized to the ER as we previously reported for these assays(26), making it difficult to spatially define donor and acceptor membranes. We therefore focused on Giantin, which remained intact under rundown conditions, providing a better view of the spatial organization of the ERES-cis Golgi interface. A dramatic and quantitative loss of overlap between Sec24C and Giantin in the juxtanuclear Golgi region was detected after the rundown (Fig. 4A, quantified in 4B). Thus, contacts between the membranes that make up the ERES hub are destabilized in rundown incubations showing transient connectivity between ERES and its target-acceptor membranes. Consistent with these data, our EM images of ERES after the rundown showed fewer, simpler tubulovesicular clusters, an increase in Sar1-induced tubulation at ERES, and the characteristic decrease in Sec31 immunogold labeling after the rundown (Fig. 1C) (3, 24, 40, 41).

**Figure 4.**
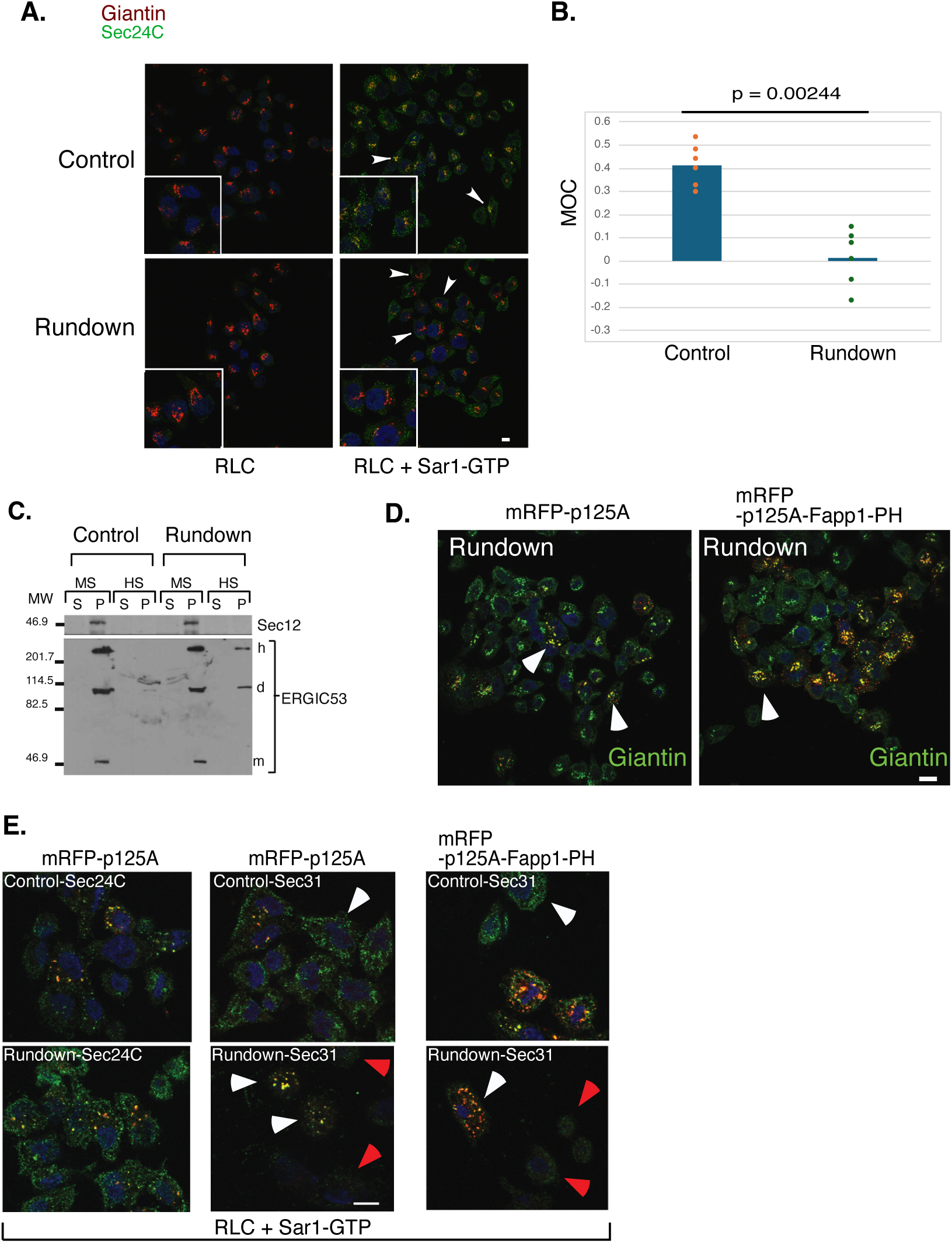
p125A-mediated PI4P recognition on acceptor membranes preserves ERES-hub architecture and COPII outer layer assembly. **A.** Permeabilized HeLa cells were incubated under control or rundown conditions (as in Fig. 1) as indicated and the assembly of Sec24C was induced by the subsequent addition of RLC and SAR1-GTP. COPII assembly at ERES (Sec24C, green) was analyzed with respect to acceptor membranes (Giantin-red). Colocalization (yellow) in merged image is shown. Arrows point to c-localization of Sec24C with Giantin in control but lack of proximity in rundown conditions. **B.** Average Manders’ overlap coefficient between Sec24C and giantin in RLC+SAR1-GTP incubations was defined using 6 cell fields, each represented by a circle (20-40 cells per field) for each condition as recorded in three independent experiments. **C.** Control or pre-incubated (rundown) microsomes (as in Fig. 2) were directly subjected to Medium Speed centrifugation (MS) and the resulting supernatant were further subjected to High-Speed centrifugation (HS) as indicated (materials and methods). Pellet (P) and supernatant (S) fractions of each step were analyzed for ERES (Sec12) and ERGIC (ERGIC53) membranes. Note the selective mobilization of ERGIC53 dimers (d) and hexamers (h) to the HS pellet under simple rundown incubation conditions. **D.** HeLa cells were transfected with mRFP-p125A or a mRFP-p125A chimera in which the DDHD domain is replaced with the FAPP1-PH domain(14) as indicated and subjected to the rundown protocol as in A, where the localization of mRFP-p125A proteins and Giantin was defined (note renewed proximity of ERES marked by p125A and Giantin and compare to panel A). Arrows point to reestablished proximity between ERES and cis Golgi in cell expressing p125A constructs as indicated. **E.** HeLa cells transiently expressing mRFP-p125A or mRFP-p125A chimera, were permeabilized and subjected to control or rundown conditions as in Fig. 1. The cells were subsequently incubated with cytosol and SAR1-GTP. The localization of mRFP-proteins (red) and the recruitment of Sec24C or Sec31 was determined by IF. White arrowheads point to ERES or rescued ERES in rundown conditions that express mRFP-p125A proteins whereas red arrowheads point to neighboring un transfected cells deficient in Sec31 assembly. Labeled nuclei (dapi, blue). Bars in A (5μm), D (20μm) and E (10μm).

### ERES-ERGIC membranes detachment in microsome-based rundown assays

If the separation of ERES-ERGIC-cis Golgi (donor-acceptor) membranes uncoupled the two COPII layers, similar membrane separation should also be observed in microsome-based assays. To determine if ER and ERGIC membranes separate during rundown, we subjected control and “rundown” microsomes to differential centrifugation and monitored changes in membrane distribution between a medium speed pellet fraction (MSP) containing heavier membranes, and high-speed pellet fraction (HSP) that contains vesiculo-tubular membranes (42). ER buds (donor membranes) were followed using mSec12 (43, 44) and ERGIC53 was used to mark ERGIC (acceptor membranes). While in control incubations both markers were confined to the MSP, in rundown incubations a fraction of ERGIC53 was found in the HSP. Interestingly, we detected mostly dimers and hexamers of ERGIC53 in this lighter fraction, which had detached from ERES in rundown conditions (Fig. 4C). Oligomeric ERGIC53 resides in a post-ER (ERGIC) compartment, as assembly takes place during ER exit (45). The observed shift of ERGIC53 from the heavy membrane fraction is not a result of budding, as under these incubations cytosolic factors were not provided. These data suggest that the biochemical analysis recapitulated our microscopy-based observations, an initial hub of connected ERES-ERGIC membranes become detached and the ERES separates from the ERGIC during the rundown. Though the microscopy-based morphological changes were more striking than the differential centrifugation outcomes, this is to be expected as there is only partial separation of donor-acceptor membrane using differential centrifugation. Overall, the two assays generated consistent results, suggesting that ERES-hub membranes detach during rundown incubations (Fig. 4A-C). The correlation between loss of membrane apposition and inhibition of coat outer layer assembly supports a model in which COPII assembly is sensitive to donor-acceptor membrane organization of ERES whereby p125A, which binds PI4P, can utilize acceptor membrane binding to support outer layer assembly at ERES (Fig. 7B).

We next examined if p125A can reverse the effects on Sec13/31 assembly at ERES in our cell-free rundown assay. p125A was over expressed in HeLa cells that were subsequently subjected to permeabilization and the rundown incubation protocol. Expressed p125A was retained in cells after permeabilization and colocalized with Sar1-recruited Sec24C at ERES both in control and rundown conditions (Fig. 4E). p125A also preserved the apposition between ERES and acceptor cis Golgi membranes under rundown conditions (Compare Fig. 4A to 4D). Importantly, p125A maintained Sec31-Sec24C association at ERES effectively reversing the rundown effect in these incubations (Fig. 4E).

Our results suggest that PI4P binding may direct p125A to bind and retain acceptor membranes while supporting COPII assembly. To examine this hypothesis, we utilized a p125A chimera in which the SAM-DDHD PI4P recognition module was replaced with a module containing an oligomerization-defective SAM domain and importantly, a bona fide PI4P-binding domain, the PH domain of FAPP1. The expression of a DDHD-truncated form of p125A in cells inhibits the assembly of endogenous Sec31/13 at ERES yet COPII assembly is restored with the expression of this p125A-FAPP1-PH chimera (14). In rundown assays, expression of the p125A-FAPP1-PH chimera protected membrane contacts at ERES, preserving the close apposition between Giantin and ERES (Fig. 4D). The chimera also preserved association of Sec31 at these sites under rundown conditions (Fig. 4E). Therefore p125A, which binds the C-terminal helical domain of Sec31A, uses PI4P recognition to preserve membrane contacts and COPII assembly under rundown conditions.

Tether and fusion proteins including Rab1, p115 or syntaxin18 regulate COPII assembly at ERES (39, 46, 47) and may support membrane apposition. However, the expression of active Rab1-GTP (Rab1a ^Q70L^) gave variable results in the cell free rundown assay. Although transfected Rab1 was retained in permeabilized cells, it did not robustly reverse the selective recruitment inhibition of the coat outer layer under rundown conditions (Sec13, Fig. S1A). Rab1 and tethers dynamically maintain ERES-ERGIC-cis Golgi contacts thus indirectly regulating coat assembly, whereas p125A allows the coat to detect these membrane contacts. Our cell free assays suggest a simple model in which COPII assembly on ER membranes utilizes p125A to interact in trans with acceptor membranes, effectively coordinating donor-acceptor membrane connectivity with coat assembly (model, Fig. 8B).

### p125A is required for the dynamic assembly of the COPII outer layer

It may be expected that expression of an outer layer associated PI4P-binding adaptor in an invitro assembly assay can protect both donor acceptor membrane contacts and outer layer assembly at ERES. Furthermore, cell-free assays utilize non-physiological stoichiometries of traffic components, and coat dynamics derived from cycles of GTP binding and hydrolysis is negated by the utilization of the Sar1-GTP mutant. We looked for an alternative physiological avenue to examine the role of p125A in directing inner-outer layer coupling. Cell free analyses predict that lack of p125A should selectively destabilize COPII outer layer recruitment at ERES. We utilized CRISPR/Cas9 in U2OS cells to delete p125A, (Fig. 5A). Deleted cells did not upregulate the expression of COPII inner (Sec23) or outer (Sec31A) layers (Fig.5A) making these cells suitable for the analysis of COPII assembly. As we previously reported with acute RNAi-mediated depletion, p125A-KO cells displayed marked effects on Golgi showing massive vesiculation and breakdown of juxtanuclear assembly (Fig. 5B-G and 5I). The effects were mirrored with various cis/medial Golgi markers, including giantin, GRASP65 and gpp130, all presenting vesiculated appearance regardless of overlap (Fig. 5B-G). We used GRASP65 to visualize the reversal of these effects by the expression of WT p125A, but not of a mutant that is deficient in PI4P binding (termed PIX), as we previously quantified in our RNAi-based studies (Fig. 5E) (14). Analysis of KO cells by transmission EM identified Golgi mini stacks, indicating that vesiculated Golgi morphology is a result of unlinking of the Golgi ribbon (Fig. 5C). These changes are typically observed in cells presenting secretion defects and cellular stress. The morphology of the coat inner layer (Sec24C) was largely unaffected by p125A deletion (Fig. 5F, SR-STED in lower panel and Fig. 5H). Hence p125A is not critical for the recruitment of the coat inner layer and its assembly at ERES. In marked contrast, Sec31A, which displayed a defined punctate appearance of ERES in parental cells, lost this defined localization showing smaller puncta and enhanced diffused appearance likely representing defects in membrane binding and increased cytosolic population (Fig. 5G-I and SR -STED in lower panels). To test for the latter, we subjected parental and p125A-KO cells to a brief permeabilization on ice and monitored the retention of the coat outer layer at ERES. Under these conditions intact Golgi assembly was perturbed over time likely due to the expected disarrayed microtubule organization. In parental cells, Sec31A remained associated with ERES following brief permeabilization while it was mostly lost from p125A-deleted cells within 5 minutes (Fig. 5I and quantified in panels below). The results suggest that p125A stabilizes Sec13/31 assembly at ERES leading to increase in membrane-bound fraction. Overall, the results provide independent validation to cell fee analyses. In permeabilized cells, expression of p125A preserves membrane organization and COPII outer layer assembly at ERES. In an orthogonal approach, deleting p125A in cells leads to perturbation of Sec31 assembly and membrane binding. Thus, our model (Fig. 7B) is supported by complimentary, independent experimental approaches.

**Figure 5.**
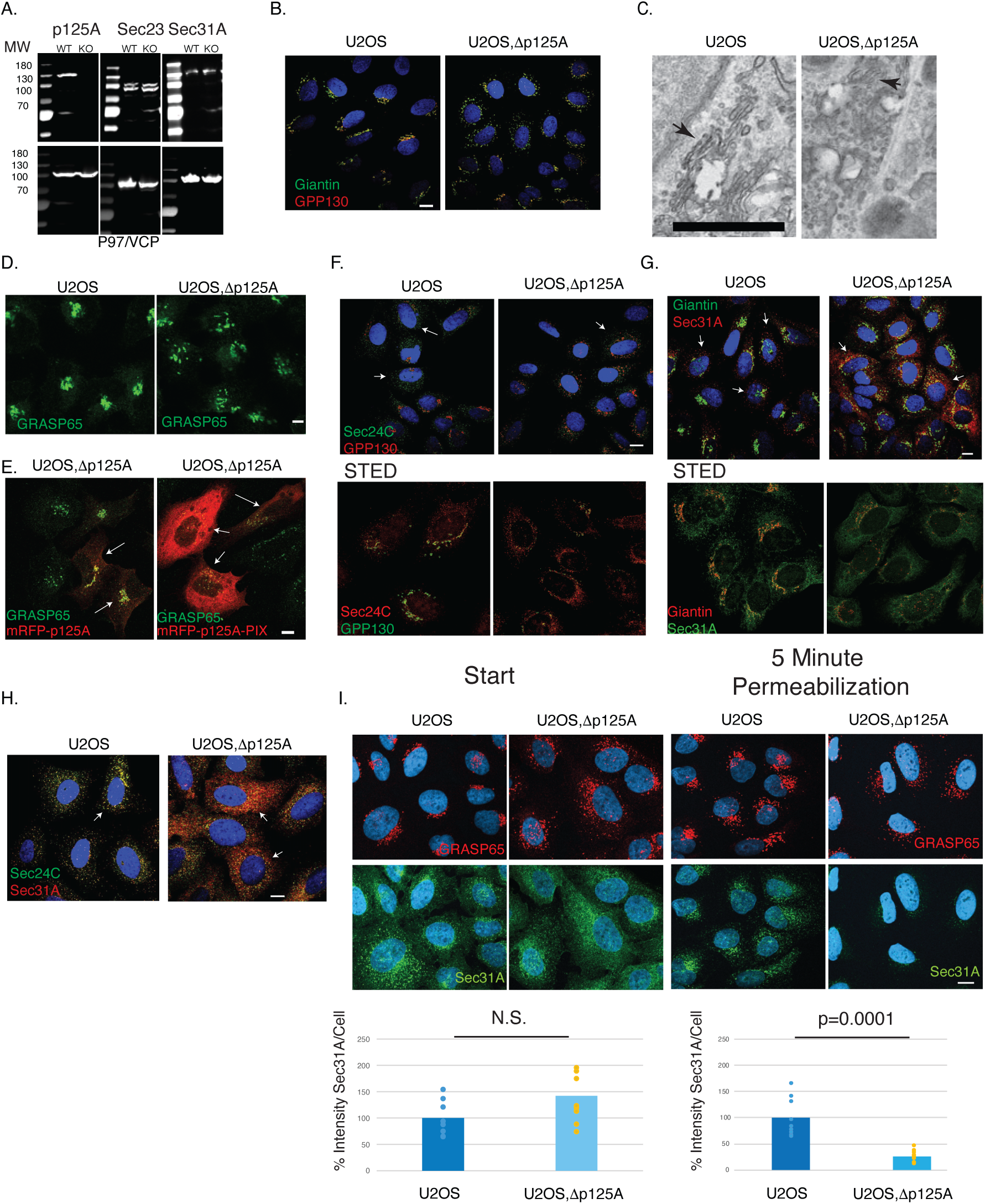
The Assembly of COPII outer layer is destabilized in p125A-KO cells. **A.** Analysis of p125A, COPII inner layer (Sec23), and outer layer (Sec31A) expression in U2OS and p125A KO cells by western blot as indicated (30 μg of protein loaded per lane, re-probed for VCP/p97 as indicated). **B.** The localization of Giantin (green) and GPP130 (red) in U2OS or U2OSΔp125A cells as indicated. **C.** Transmission EM of U2OS or U2OSΔp125A cells showing stacked Golgi elements (arrows). Bar is 1μm. **D.** The localization of GRASP65 in U2OS or U2OSΔp125A cells. **E.** U2OS or U2OSΔp125A were transfected with mRFP-p125A or p125A-PIX mutant were analyzed for GRASP65 localization. Arrows point to transfected cells showing variable expression levels. **F.** The localization of COPII inner layer (Sec24C, green for confocal image, red for STED, lower panel) and cis Golgi (GPP130, red in confocal, green in STED) in U2OS or U2OSΔp125A. Arrows point to Sec24C coated ERES. **G.** The localization of COPII outer layer (Sec31A, red in confocal, green in STED, lower panel) and cis Golgi (Giantin, green in confocal, red in STED) in U2OS or U2OSΔp125A. Arrows point to Sec31A coated ERES and lack of localization in U2OSΔp125A cells **H.** The localization of COPII outer (Sec31A, red) and inner (Sec24C, green) layers in U2OS or U2OSΔp125A cells. **I.** Parental U2OS or U2OSΔp125A cells were stained with Sec31A and GRASP65 antibodies or permeabilized with saponin for 5 min on ice, fixed and stained for GRASP65 and Sec31A. Lower panels, quantification of Sec31A intensity per cell in parental and U2OSΔp125A cells, in start (left) and 5 min. permeabilization on ice (right) Circles represent random cell fields (15-45 cells) derived from two independent experiments, normalized to parental cells (100%) and analyzed using two-tailed t-test. Bars for IF are 10μm.

### Reprograming of ECM Remodeling and Secretion in p125A-Deleted Cells

Deleting p125A strongly affects COPII outer layer assembly, so we examined how cells adapt to maintain function. We compared the gene expression (transcriptome) of control U2OS and p125A-deleted U2OS cells. While cells of the same group showed little variation (Fig. S2A), p125A-knockout cells had considerable changes (Figs. 6A, S2 and S3), with 1115 genes upregulated and 1252 downregulated (Fig. 6B). Gene Ontology (GO) analysis showed that pathways related to extracellular matrix (ECM) remodeling were most affected, including ECM structure, collagen organization, and metabolism (Fig. 6C). Similarly, many downregulated genes also belonged to ECM-related categories and pathways involved in ECM interactions, remodeling, organization and ones directing collagen biosynthesis, trimerization and fibril assembly were overrepresented (Figs. S3 and S4), indicating a comprehensive and coordinated shift in gene expression to adjust ECM composition and function in response to p125A loss.

**Figure 6.**
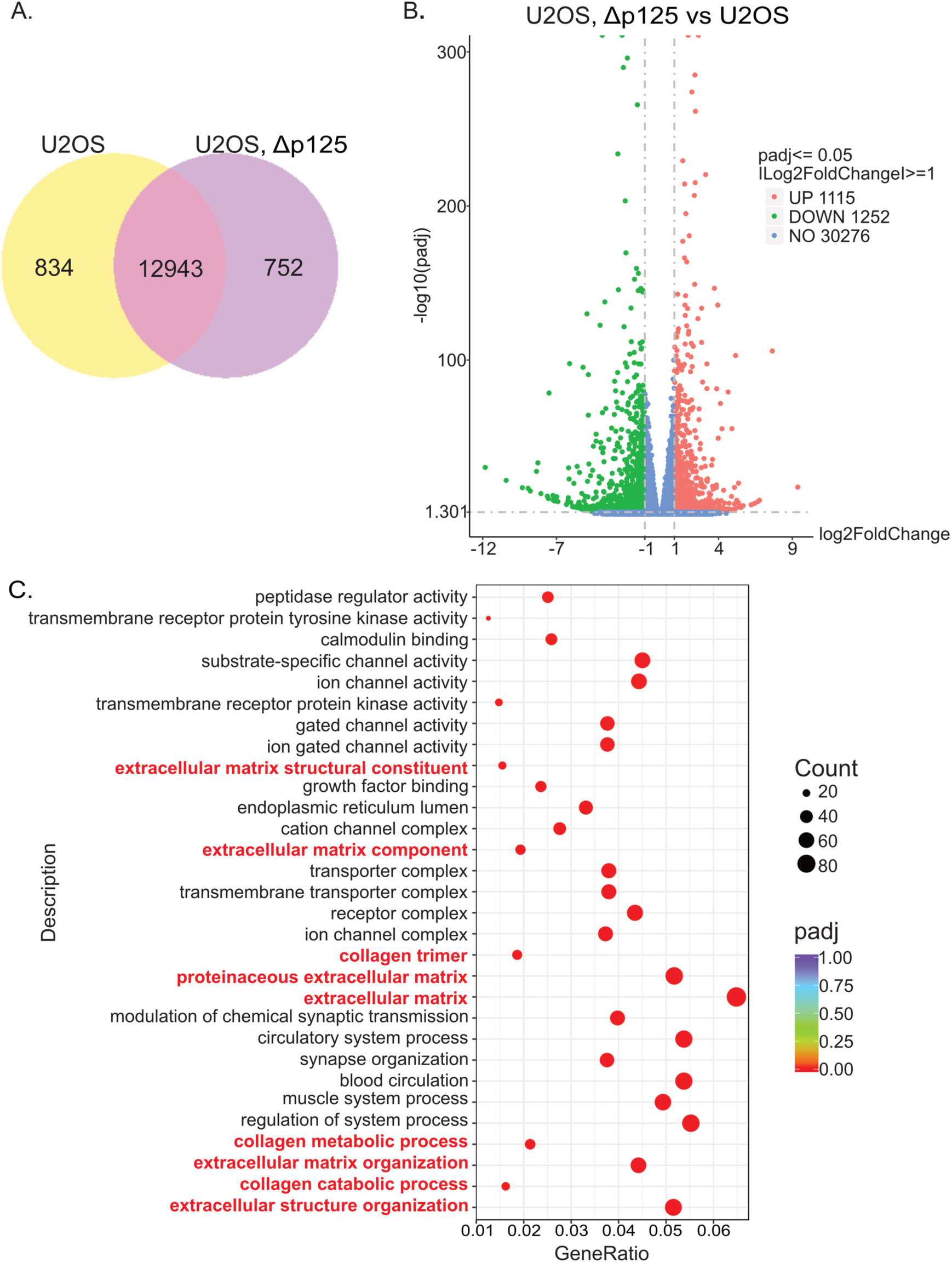
Transcriptomic analysis of U2OSΔp125 and U2OS. **A.** The co-expression Venn diagram presents the number of genes that are uniquely expressed within each group, with the overlapping region showing the number of genes that are co-expressed in both U2OS and U2OSΔp125. **B.** Volcano plot of differentially expressed genes (DEGs) (padj < 0.05 and |log 2 (FoldChange)| > 1) identified between U2OSΔp125 and U2OS. The x-axis represents the log 2 (FoldChange), y-axis represents statistical significance for each gene. Green dots denote down-regulated gene expression, red dots denote up-regulated gene expression in U2OSΔp125. The blue dots denote gene with unchanged expression. The horizontal dotted line indicates the padj-value cut-off, while the vertical dotted lines mark the point for no change.**C.** Dot plot of the Gene Ontology (GO) enrichment analysis for DEGs. The most significant 30 GO Terms were selected for display. The x-axis is the ratio of the number of differential genes linked with the GO Term to the total number of differential genes, and the y-axis is GO Terms. The size of a point represents the number of genes annotated to a specific GO Term, and the colour from red to purple represents the significant level of the enrichment. GO pathways with padj < 0.05 are significant enrichment. (See Supplementary Table 1 for complete gene lists).

To explore how cells compensate, we analyzed protein secretion using mass spectrometry (Table 1, Supplemental Tables 2-4). The secretion of many ECM components and modifying enzymes, such as matrix metalloproteinase 9, Matrilin-3, and lysyl oxidase homologs 2 and 4, was inhibited in p125A-knockout cells (Table 1). Consistently, secretion of fibrillar collagens, including COL1A1, was reduced (Table 1). However, a few proteins, like collagen XVI, a non-fibrillar FACIT (fibril-associated collagens with interrupted triple helices) collagen, were hypersecreted (Supplement Table 3). These findings suggest that cells may adjust the ECM composition to compensate for loss of p125A and the potential derived defects in biosynthetic secretion of ECM components including bulky fibrillar cargoes.

**Table 1.**
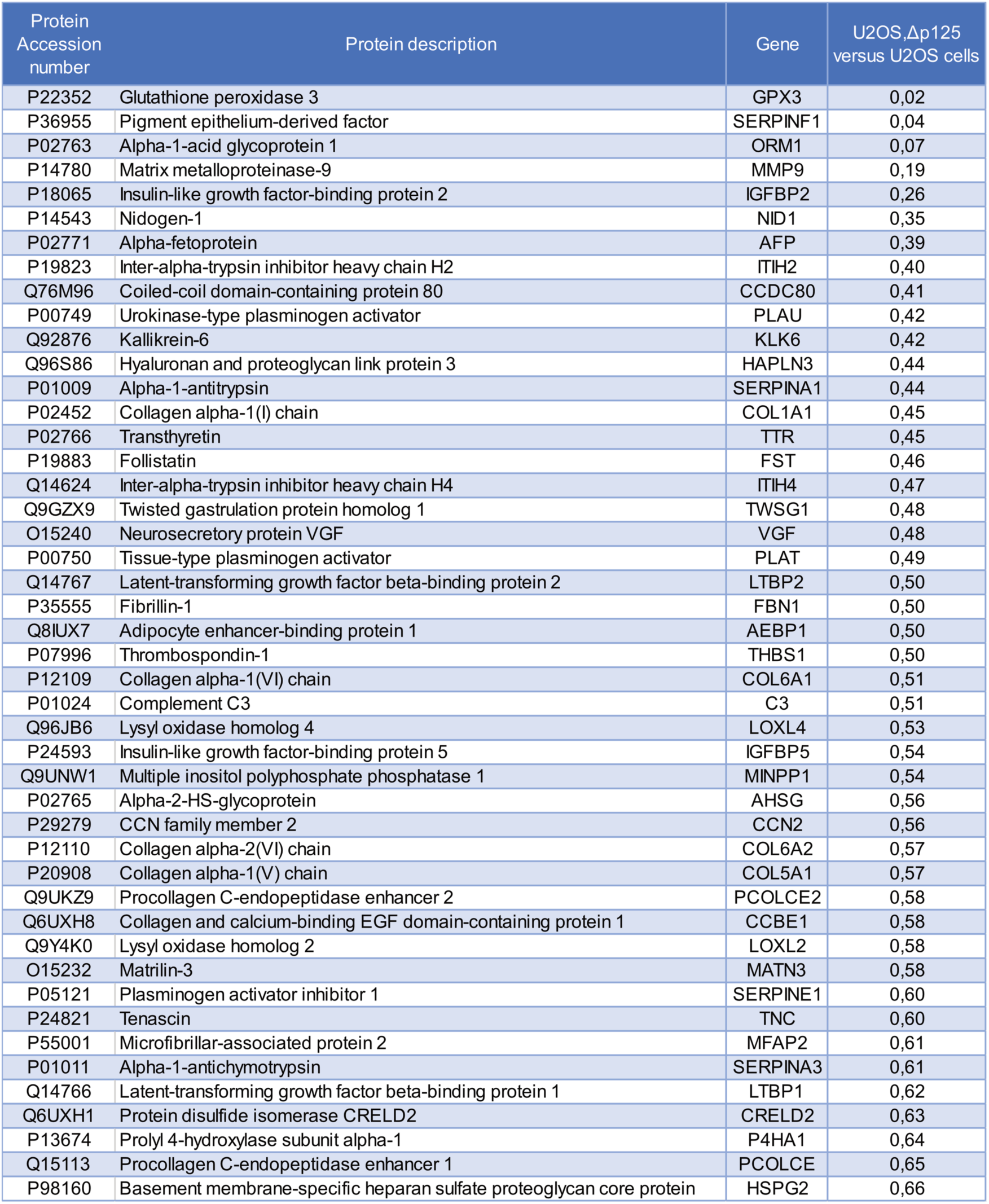
List of proteins identified as hypo-secreted in U2OSΔp125 compared to U2OS. Secreted and intracellular proteins were quantified by LC-MS/MS in U2OSΔp125 and U2OS cells. Within each group, the proportion of the secreted protein was determined. Then, each value for the U2OSΔp125 was compared to the control U2OS. 46 proteins were finally identified as less secreted with a threshold set at 0.67. (See Supplementary Table 4 for complete proteins lists detected in lysate and secretome).

| Protein Accession number | Protein description | Gene | U2OS,Δp125 versus U2OS cells |
| --- | --- | --- | --- |
| P22352 | Glutathione peroxidase 3 | GPX3 | 0,02 |
| P36955 | Pigment epithelium-derived factor | SERPINF1 | 0,04 |
| P02763 | Alpha-1-acid glycoprotein 1 | ORM1 | 0,07 |
| P14780 | Matrix metalloproteinase-9 | MMP9 | 0,19 |
| P18065 | Insulin-like growth factor-binding protein 2 | IGFBP2 | 0,26 |
| P14543 | Nidogen-1 | NID1 | 0,35 |
| P02771 | Alpha-fetoprotein | AFP | 0,39 |
| P19823 | Inter-alpha-trypsin inhibitor heavy chain H2 | ITIH2 | 0,40 |
| Q76M96 | Coiled-coil domain-containing protein 80 | CCDC80 | 0,41 |
| P00749 | Urokinase-type plasminogen activator | PLAU | 0,42 |
| Q92876 | Kallikrein-6 | KLK6 | 0,42 |
| Q96S86 | Hyaluronan and proteoglycan link protein 3 | HAPLN3 | 0,44 |
| P01009 | Alpha-1-antitrypsin | SERPINA1 | 0,44 |
| P02452 | Collagen alpha-1(I) chain | COL1A1 | 0,45 |
| P02766 | Transthyretin | TTR | 0,45 |
| P19883 | Follistatin | FST | 0,46 |
| Q14624 | Inter-alpha-trypsin inhibitor heavy chain H4 | ITIH4 | 0,47 |
| Q9GZX9 | Twisted gastrulation protein homolog 1 | TWSG1 | 0,48 |
| O15240 | Neurosecretory protein VGF | VGF | 0,48 |
| P00750 | Tissue-type plasminogen activator | PLAT | 0,49 |
| Q14767 | Latent-transforming growth factor beta-binding protein 2 | LTBP2 | 0,50 |
| P35555 | Fibrillin-1 | FBN1 | 0,50 |
| Q8IUX7 | Adipocyte enhancer-binding protein 1 | AEBP1 | 0,50 |
| P07996 | Thrombospondin-1 | THBS1 | 0,50 |
| P12109 | Collagen alpha-1(VI) chain | COL6A1 | 0,51 |
| P01024 | Complement C3 | C3 | 0,51 |
| Q96JB6 | Lysyl oxidase homolog 4 | LOXL4 | 0,53 |
| P24593 | Insulin-like growth factor-binding protein 5 | IGFBP5 | 0,54 |
| Q9UNW1 | Multiple inositol polyphosphate phosphatase 1 | MINPP1 | 0,54 |
| P02765 | Alpha-2-HS-glycoprotein | AHSG | 0,56 |
| P29279 | CCN family member 2 | CCN2 | 0,56 |
| P12110 | Collagen alpha-2(VI) chain | COL6A2 | 0,57 |
| P20908 | Collagen alpha-1(V) chain | COL5A1 | 0,57 |
| Q9UKZ9 | Procollagen C-endopeptidase enhancer 2 | PCOLCE2 | 0,58 |
| Q6UXH8 | Collagen and calcium-binding EGF domain-containing protein 1 | CCBE1 | 0,58 |
| Q9Y4K0 | Lysyl oxidase homolog 2 | LOXL2 | 0,58 |
| O15232 | Matrilin-3 | MATN3 | 0,58 |
| P05121 | Plasminogen activator inhibitor 1 | SERPINE1 | 0,60 |
| P24821 | Tenascin | TNC | 0,60 |
| P55001 | Microfibrillar-associated protein 2 | MFAP2 | 0,61 |
| P01011 | Alpha-1-antichymotrypsin | SERPINA3 | 0,61 |
| Q14766 | Latent-transforming growth factor beta-binding protein 1 | LTBP1 | 0,62 |
| Q6UXH1 | Protein disulfide isomerase CRELD2 | CRELD2 | 0,63 |
| P13674 | Prolyl 4-hydroxylase subunit alpha-1 | P4HA1 | 0,64 |
| Q15113 | Procollagen C-endopeptidase enhancer 1 | PCOLCE | 0,65 |
| P98160 | Basement membrane-specific heparan sulfate proteoglycan core protein | HSPG2 | 0,66 |

### Spatial sensing by p125A selectively coordinates COPII with collagen traffic

Fibrillar procollagens are too large for COPII vesicles, and both imaging and functional analyses demonstrate that such bulky cargoes are transported via tunnels between ERES and ERGIC membranes (48). We examined if a lack of p125A inhibits procollagen 1 secretion from the ER. At steady state, a minimal amount of intracellular fibrillar procollagen (COL1A1) was detected in parental U2OS cells, mainly localizing to the juxtanuclear Golgi region with occasional cells showing more accumulation. In contrast, many of the p125A-KO cells accumulated procollagen in expanded beaded ER-like morphology (Fig. 7A, quantification in lower panel) in agreement with the mass spectrometry analyses (table 1).

**Figure 7.**
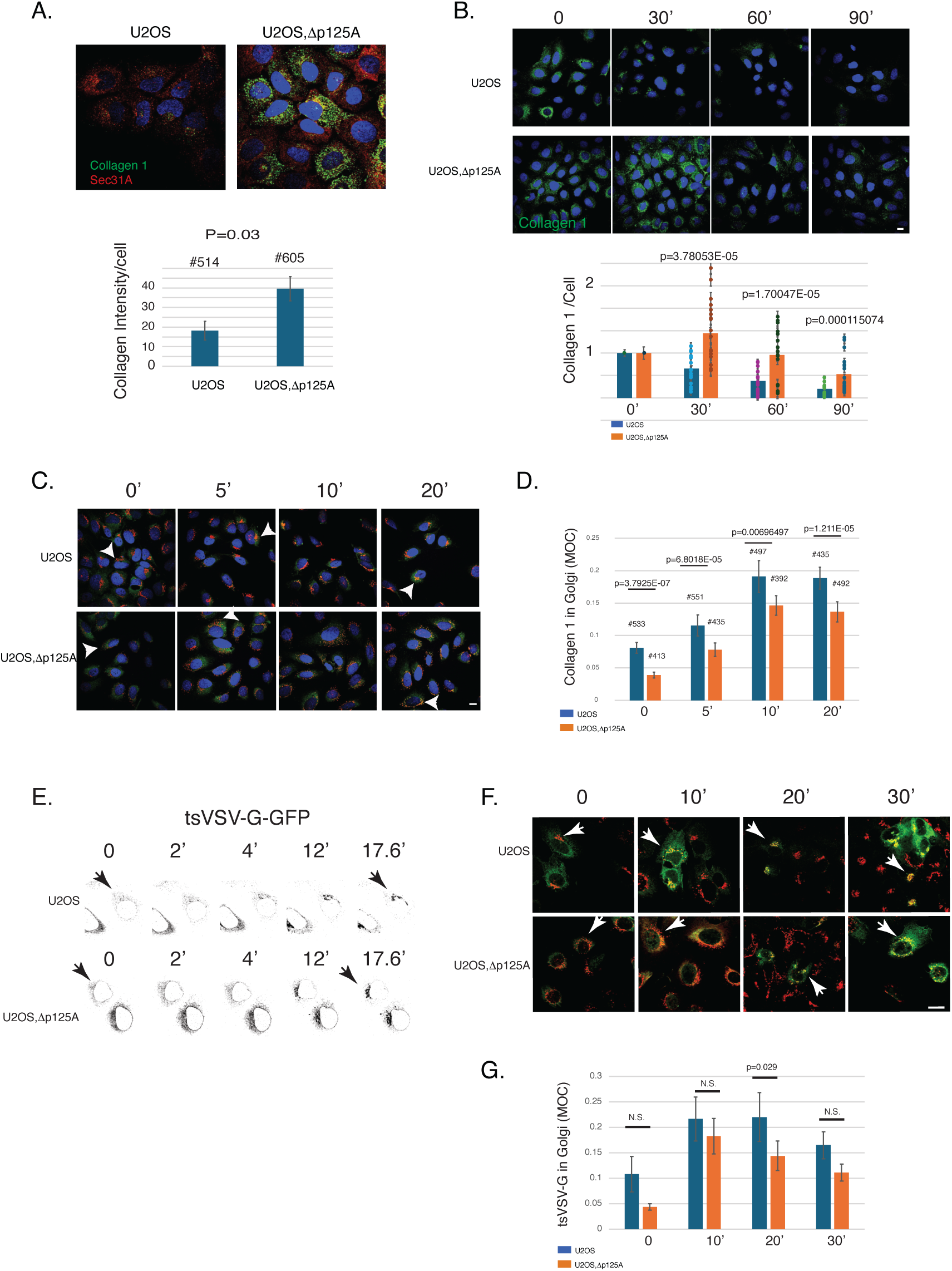
Biosynthetic traffic in p125A KO cells. **A.** The distribution of Sec31A (red) and Collagen 1 (green) in U2OS or U2OSΔp125A as indicated. Lower panel, total intensity of collagen in U2OS or U2OSΔp125A, using 2 independent samples for each quantifying 4 fields of cells per sample (two-tail T-test between groups and the total number of cells quantified per bar are shown with SEM). **B.** U2OS or U2OSΔp125A were incubated at 40°C for 3 hr. in the presence of 1% FCS. Following incubations, ascorbate and cycloheximide were added and the cells were shifted to 32°C for the indicated time to monitor Collagen 1 levels. Lower panel: quantification of remaining collagen 1 levels per cell for each time point as indicated, normalized to time 0. Each circle represents a field with an average of 22 cells. Data are taken from four independent experiments and shown with two tailed T-test between groups and SEM. **C.** U2OS or U2OSΔp125A were incubated as in B for the indicated time points and stained for collagen 1 (green) and GPP130 (red). **D.** Quantification of co-localization (Manders’ Overlap Coefficient, MOC) between collagen 1 and GPP130 in U2OS or U2OSΔp125A cells at the indicated time points. Shown are averages from 4 independent experiments where at least 3 fields of cells (∼10 cells/field) were quantified per time point per each experiment (with two tail t-test and total number of cells quantified per bar, shown with SEM). **E.** U2OS or U2OSΔp125A as indicated, accumulating transfected tsVSV-G-GFP in the ER at 40°C were shifted to 32°C in the presence of cycloheximide and live-imaging was carried. Indicated time points in minutes, movies in supplement. **F.** U2OS or U2OSΔp125A were shifted to 32°C and incubated as in E, fixed at indicated time points and stained for tsVSV-G-GFP (green) and GPP130 (red). **G.** Quantification of tsVSV-G-GFP and GPP130 overlap (Manders’ Overlap Coefficient, MOC) in U2OS or U2OSΔp125A cells at indicated times, averaged from 5 independent experiments where at least 3 fields of transfected cells per experiment were quantified per time point (two tail t-test for each time point shown with SEM). Arrows in panels C, E and F point to the Cis Golgi region. Scale bars are 10μm.

To examine for potential traffic defects that may account for higher collagen 1 levels in KO cells, we subjected parental and p125A KO U2OS cells to increased temperature (40°C) and ascorbic acid depletion. These treatments interfere with procollagen prolyl hydroxylation and triple helical assembly, leading to accumulation in the ER. Subsequently, cycloheximide and ascorbate were added, and cells were incubated at 32°C to initiate traffic. We monitored the clearance of collagen 1 from parental and p125A-KO cells over 90 minutes. Levels of collagen 1 were markedly decreased over the incubation period in parental cells, yet secretion was markedly delayed in KO cells retaining collagen 1 from up to an hour before the beginning of clearance (Fig. 7B, quantification in lower panel). Therefore p125A-KO present a clear defect in collagen 1 secretion.

To examine traffic from ERES, where p125A-COPII localizes and functions, we analyzed the traffic of collagen 1 to the Golgi complex by quantifying overall co-localization of collagen with a Golgi marker (gpp130) over time using Mander’s overlap coefficient. In WT cells procollagen 1 accumulated in the ER began to clear, and increased overlap was observed in the Golgi within 5-10 minutes. In KO cells procollagen 1 mobilization as reflected by Golgi localization was significantly reduced throughout the time course of this analysis (Fig. 7C and quantification in 7D). Therefore, the overall secretion defect of collagen 1 (Fig. 1A-B) is mirrored by a reduced localization of collagen 1 in the Golgi at the initiation of secretion.

The defect in collagen secretion may represent a global defect in traffic from the ER. However, secretome analysis does not support a global effect (Table 1 and supplemental tables 2-3). Alternatively, the inability to coordinate COPII layer-linkage with donor-acceptor contacts required for tunnel-based traffic may selectively affect the mobilization of bulky cargo. To explore these possibilities, we utilized the temperature sensitive Vesicular Stomatitis Virus G protein (tsVSV-G), an established reporter for the traffic of small size cargoes for analysis. tsVSV-G accumulates in the ER at a non-permissive temperature (40°C) and effectively mobilizes out of the ER in a synchronous manner when shifted to a permissive temperature (32°C). We followed a GFP-tagged tsVSV-G using live imaging. Upon a temperature shift, the reporter which resided in the reticulum concentrated and mobilized from the ER to the perinuclear region effectively in both WT and KO cells, with an overall similar kinetics (Fig. 7E and supplement movies 1 and 2). To quantify ER-Golgi traffic, we monitored the overlap of tsVSV-G with cis Golgi at select time points in fixed cells. Overall, tsVSV-G overlap with cis Golgi was similarly increased in both WT and KO cells suggesting that the traffic defect in p125A deleted cells is selective for collagen (Fig. 7F-G). The results agree with our secretome analysis (tables 1 and S3) and collectively suggest that when spatial sensing of donor-acceptor membrane apposition is perturbed, the coordination between coat assembly and traffic is compromised, leading to selective defects in mobilization of cargo that requires direct non vesicular traffic.

## Discussion

The vesicular traffic model offers an elegant solution for preserving compartment identities while facilitating efficient transport of approximately a third of the human proteome between cellular compartments and to various cellular and extracellular destinations. The model has contributed to the identification of vesicle forming coats, crucial components utilized in intracellular traffic(49). However, a vesicle-centric traffic model falls short of addressing the transport of significant cargoes including proteolipid particles and ECM components, particularly fibrillar collagens, that are synthesized in the ER. These cargoes, which include collagens, account for about 20% of the human dry body weigh yet do not conform to the size constraints of COPII vesicles involved in traffic.

Accumulating data produced strong evidence for a role of tunnels in supporting general secretion and specifically as an added key mode of traffic from the ER(2), raising a fundamental question: How can vesicle-forming coats function in non-vesicular, tunnel-based traffic. In this work, we address this question by providing evidence that COPII coat assembly is coordinated with donor-acceptor membrane proximity required for connectivity and tunnel-based ER export (Figs 1-4). We provide evidence that p125A functions as a COPII coat adaptor that couples coat-assembly with ERES-ERGIC/cis Golgi organization (Figs 4-5) and is selectively required for tunnel-based traffic (Figs. 6-7).

Functional and morphological evidence support direct tunnel-based traffic from the ER, where ERGIC compartments become connected to ERES (2, 9). The morphology of ER exit tunnels suggests that coordination between COPII-vesicle budding-fission and subsequent tethering and fusion generates tunnels composed of chain of 1 to 3 connected vesicles (Fig. 8A). This hints at a potential inclination towards traffic route influenced by cargo load. The equilibrium is tilted toward vesicular traffic in drosophila yet prefers tunnels in mammalian cells (9, 38). The formation of tunnels may alternatively be driven by retrograde traffic as implied from the utilization of the NRZ tether and retrograde fusion proteins in the exit of procollagen from the ER and from the ability to maintain the traffic of small sized cargoes under conditions where COPII is depleted(11, 50). Both models are not mutually exclusive. Regardless of mechanisms that build tunnels, our cell-free assays suggest that building of tunnels between compartments is a dynamic process involving transient connections between donor and acceptor membrane compartments (Figs. 1E and 4). In cell-free conditions, donor-acceptor membrane detachment becomes irreversible due to the imposed (and relatively infinite) dilution of the reactions, leading to an observed *rundown* effect (Figs. 1E and 4). Overall, the assays likely recapitulate a single step in an otherwise dynamic cycles of tethering and fusion that support exit of cargoes from the ER (48, 51).

**Figure 8.**
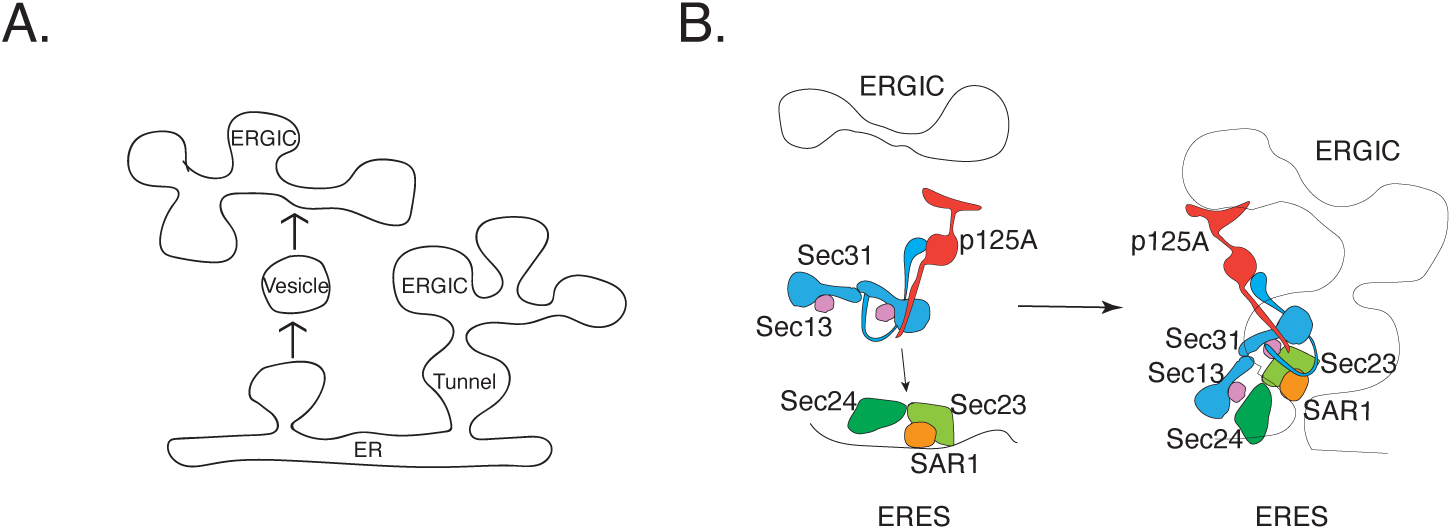
**A.** Model describing vesicular vs. tunnel-based traffic. **B.** Model describing spatial sensing by the COPII coat. 1. The cytosolic COPII inner layer (SAR1-Sec23/24) is recruited onto donor ERES membranes (left panel). The cytosolic COPII outer layer, composed of Sec13/31 and its associated adaptor p125A (Sec23ip) is recruited onto SAR1-Sec23/24 while p125A binds Sec23. Sec13/31 recruitment onto the coat inner layer is stabilized by selective interactions between p125A and PI4P lipid signals that are enriched on ERGIC acceptor membranes (right panel).

### p125A connects COPII assembly with the recruitment of ERGIC/cis Golgi to the ERES

The mechanisms by which the balance between tunnel or vesicle formation at ERES is executed is unknown. ERES Sec12-cTAGE5-TANGO1-Sec16 fence complexes interact with COPII to facilitate inner layer trapping and Sar1 activation while delaying outer layer assembly (33). The TANGO1 fence further facilitates ERES-ERGIC-cis Golgi membrane recruitment (12). The ability to regulate coat assembly while driving membrane contacts support a role for the TANGO1-cTAGE5 fence in controlling the balance between vesicles and tunnels(38, 52). TANGO1 deletion in drosophila and mammalian cells leads to excessive vesiculation that inhibits overall secretion of a variety of cargoes including that of a small size bulk flow marker that accumulate in the ER at steady state(53). Thus donor-acceptor transient connectivity is critical for effective physiological biosynthetic secretion. However, the deletion of p125A does not inhibit general secretion. Similar outcomes are observed when the p125A binding COPII outer layer is depleted, or in general when COPII functions are compromised leading to selective inhibition of collagen secretion(24, 34, 50, 54–56). A tight nanometer-scale interface between ERES-ERGIC likely allows for processes of coat mediated budding, tether and fusion to proceed even when coat assembly is uncoordinated. In fact, such processes proceed even when COPII levels are severely depleted(50), yet the coordination of coat assembly with tether-mediated contacts and fusion becomes significant for bulky cargoes relying on tunnel-based traffic (Fig. 7).

Our results suggest that ERES-ERGIC-cis Golgi membrane contacts are transmitted to the coat outer layer to coordinate coat assembly that facilitates fission. This is achieved by using a key adaptor that is stoichiometric component of the coat outer layer-p125A. p125A selectively binds to acceptor membranes enriched in PI4P (ERGIC and/or cis-Golgi), while controlling COPII coat assembly at the ERES by linking the two coat layers (Fig. 5 and model in 8B). The cell free coat assembly assays rely on negating Sar1 dynamics locking it in a GTP bound state. The fact that a differential stabilization of the two coat layers is observed in our cell free assays (Figs. 1-2) agrees with previous data showing that expression of a GTPase deficient (Sar1-GTP) in cells does not negate the dynamic exchange of the coat outer layer (Sec13) at ERES (57). Our result show that the linkage between inner and outer layers is governed by added interactions, including the recognition of acceptor membranes by p125A or by a chimera of p125A harboring a bona-fide PI4P-binding domain.

The regulation of inner-outer layer coupling is conserved in evolution. The ERES scaffold protein Sec16p controls inner-outer coat assembly in budding yeast, regulating traffic (29). This regulation is expanded in evolution with added dedicated adaptors that control inner-outer layer coupling to couple physiological needs with coat activities. CREB regulated transcription coactivator 2 (CRTC2) modulates COPII-dependent SREBP1 processing by competing with Sec23A for Sec31A binding (58). COPII adaptors that respond to signaling cues such as elevated Ca^2+^ concentration via proteins like Alg2, or selective lipid signals via p125A, further control coat inner-outer layer coupling. These adaptors interact with different domains on the C-terminus of Sec31, with Alg2 binding the proline rich linker and p125A interacting with the structured-terminal helical domain of Sec31A respectively (Fig. 3D-E). While cryoET reconstruction of COPII assembly on ER membranes in yeast presents a clear inner-outer layer topology(59), both Alg2 and p125A link the mammalian COPII outer layer with membranes and these activities are required for assembly at ERES(60). In the case of p125A lipid-recognition specificizes acceptor membranes, the role of Alg2 in membrane binding is yet undefined. The C-terminal helical domain, which is required for Sec31p functionality and yeast viability(61), directs proper assembly of the outer layer at mammalian ERES (Fig. 3A-C). Over expressed p125A phase separates at ERES and collects acceptor compartments and this activity requires the Sec23-binding N-terminal IDR and C-terminal PI4P binding(14). We find that expression of p125A or a chimera in which the lipid binding module is replaced with an otherwise Golgi localized PI4P binding domain (Fapp1-PH) rescues the assembly of COPII outer layer in cells (14) and in rundown-based cell free assays (Fig. 4) (15, 62).

### p125A is required for collagen traffic

The destabilization of outer layer assembly observed in p125A deleted cells leads to inhibition of COL1A1 traffic (Figs. 6-7). The significance of this regulation is evident in humans, where a homozygous patient expressing a DDHD-truncated form of p125A, which cannot bind PI4P, exhibits severe phenotypes reminiscent of COPII mutations that reduce neural crest cell migration and inhibit collagen secretion from the ER (63).

Our results suggest that p125A assists in spatial sensing reporting on donor acceptor apposition, which is likely maintained by TANGO1 and cTAGE5 proteins that recruit acceptor membranes to ERES. Yet in our cell free assays, p125A expression preserved membrane apposition in rundown conditions (Fig. 4D). While these results may be derived from p125A over expression, a recent study characterized an interaction between Golgi localized VPS13B and p125A that contribute to the apposition between ERES and cis Golgi membranes. Mutations in VPS13B that affect p125A binding lead to the development of Cohen syndrome, which is characterized, among other, by symptoms indicative of collagen secretion defects. VPS13B binds the N-terminus IDR of p125A whereas its C-terminal PI4P-binding module comprised of a SAM and DDHD domains is independently targeted to Golgi membranes(14). In agreement with our results, the authors find that acute depletion of p125A using RNAi leads to robust inhibition of collagen 1 secretion, and this inhibition is more pronounced than the inhibition of collagen secretion derived from VPS13B depletion(64). The combined results support our model, suggesting that when coat assembly and tunnel formation are not coordinated, collagen does not exit the ER effectively. VPS13B likely acts as a lipid tunnel, helping lipids move between ERES and Golgi membranes. If this process is passive, lipids would flow from Golgi membrane to ER-ERES, reducing membrane tension at ERES to support membrane expansion and tubulation(65).The interactions between p125A-VPS13B are specifically found at perinuclear ERES close to the cis Golgi. Our recent findings show that perinuclear ERES comprising ∼40-50% of the total ERES population are selectively utilized to mobilize collagens, suggesting that localized reduction in membrane tension may assist in collagen secretion from the ER(66). VPS13B is a minor component in cells, is not detected at peripheral ERES together with p125A and does not compete with Sec31A for p125A binding(64). Its lower expression compared with p125A (and COPII), could mean it fine-tunes lipid composition at select ERES, while p125A facilitates coat assembly across all ERES. Although p125A has no PLA1 enzyme activity when measured in isolation, such activity may be regulated locally at ERES. Whether a p125A enzyme assists in directing lipid flow to and from ERES is a topic for future research.

### COPII, a coat and a collar on transport intermediates

Our results provide an elegant, simple mechanism by which COPII can be coordinated to support traffic by detecting donor-acceptor membrane contacts and potentially tunnel formation. While it is likely that tunnels can form in the absence of COPII activity(50), the role of COPII in tunnel-based traffic requires further studies. Our work defined a regulated step that depends on membrane apposition: the coupling between the coat inner and outer layers (Fig.7). This coupling controls the rate of GTP hydrolysis by Sar1. Controlling Sar1-GTPase activation may direct the membrane deforming activities of Sar1-COPII at ERES and participate in coordinating tether and fusion activities (tunnel formation) with Sar1-GTPase-driven fission of traffic intermediates (2, 9, 67). Sar1-GTPase activation may also direct a processive concentration of a select group of cargoes at ERES for traffic (10, 68). Similarly, this regulation may govern the exclusion of ER resident proteins, providing dynamic gating at ERES required to prevent nonspecific content mixing between donor and acceptor membranes(69). The assembly of quasi-rigid hedral cages by Sec13/31 at the donor-acceptor membrane interface may also stabilize membrane connectivity, coupling coat mediated cargo sorting with direct, tunnel-based traffic(70). The utilization of phosphoinositide signals to direct COPII activities may further control other traffic routes from ERES. Future studies should address how PIP signals are utilized to direct COPII to biosynthetic secretion or autophagy/lysosomal degradation pathways.

Hub-like clustering of donor-acceptor membranes characterizes many cellular sorting sites that function in the secretory and endocytic pathways where coat activities can be similarly regulated to support vesicular and non-vesicular traffic.

## Materials and Methods

Cells. HeLa or U2OS cells were maintained at sub-confluence in Dulbecco’s Modified Eagle’s Media (DMEM; HyClone Fisher-Scientific or Lonza. Cat. N°: BE12-604F/U1) supplemented with up to 10 % Fetal Bovine Serum (FBS; Serum Source International, Inc.) and 5% Penicillin-Streptomycin (Cellgro) under standard incubation environment (37°C, 5% CO2). Transfection of HeLa cells was carried out using Lipofectamine 2000 reagent (Invitrogen-Life Technologies) according to provided protocol, with optimized DNA concentrations. Transfection of U2OS cells was carried out using X-tremeGENE™ ROCHE transfection reagent (Millipore Sigma) according to provided protocol, with optimized DNA concentrations. HeLa derived ER microsomes and rat liver cytosol were prepared as described (42).

### Plasmids

A construct for rat ECFP-RnSec31A expression was kindly provided by Dr. R. Pepperkok (EMBL Heidelberg, Germany). Stop codons at residues Q884 and I959 of ECFP-RnSec31a were generated by site-directed mutagenesis [RnSec31Q884*_F: 5’CCACAGGGGCTGTAGCAACAGCCTTCA3’; RnSec31Q884*_R: 5’TGAAGGCTGTTGCTACAGCCCCTGTGG3’; RnSec31I959*_F: AAAATTACCAAGAAGCCTTAACCAGACGAGCACCTCATT3’; RnSec31I959*_R: AATGAGGTGCTCGTCTGGTTAAGGCTTCTTGGTAATTTT3’].

The GST-fusion with human HsSec31a1041-1220 was generated by restriction enzyme cloning of HsSec31A PCR fragment into pGEX 4T-1. [HsSec31a_EcoRI_1041_F: 5’TCCCCGGAATTCTCAGCTCCAGTACCACTGTCAAGC3’; HsSec31a_XhoI_1220_R: 5’CGGCCGCTCGAGTTAGACACCCAGCTTATTGGCCTGGGT3’]. The GST-fusion proteinHsSec31a1041-1113 was generated by site-directed mutagenesis of residue E114 of GST-HsSec31a1041-1220 to a stop codon [HsSec31a_E1114*_F: 5’CCTATTCCAGATTAGCACCTCATTCTA3’; HsSec31a_E1114*_F: 5’TAGAATGAGGTGCTAATCTGGAATAGG3’].

The construct for the expression of Flag-Rab1a-GTP(Q70L) was kindly provided by Dr. Dan Devor (University of Pittsburgh). All constructs expressing FP-tagged p125A proteins were previously described (14). tsVSV-G-GFP was kindely provided by Dr. Jennifer Lippincott-Schwartz (Janelia Farms, HHMI)(71). CD25 (IL2-R alpha, TAC) fused to gp25l C-terminus provided by Dr. Linton M. Traub (University of Pittsburgh). All constructs were verified by sequencing.

### siRNA

HeLa cells were seeded at 3×105 cells/35mm dish. The next morning, cells were transfected with siRNAs to Control (DS NC1; IDT),

Sec16L 5’ CCAGGUGUUUAAGUUCAUCUACUCC-3’ and

antisense 3’ AAGGUCCACAAAUUCAAGUAGAUGAGG 5’,

or Sec31 [Sense:5’GGACAGCUUCUCUUCCACUCAATA3’;

Antisense:5’UAUUGAGUGGAAGAGAAGCUGUCCUU3’ using Lipofectamine RNAiMAX (Invitrogen-Life Technologies) according to manufacturer’s directions. siRNA transfections were repeated at 24 hours post-transfection. At 36 hours, each reaction was split onto glass coverslips in 6-well tissue culture plates. At 60 hours, cells were transfected with pECFP-rSec31a, pECFP-rSec31aQ884*, or pEYFP-Sec31aI959 with Lipofectamine 2000 (Invitrogen-Life Technologies) according to manufacturer’s directions. At 72 hours, cells were fixed for direct immunofluorescence or lysed for WB analysis with p125A, Sec31a or p97/VCP ATPase-specific antibodies.

### Rundown Biochemical Recruitment

The recruitment of Sec23 and Sec31 to ER microsomes prepared from HeLa cells was performed as previously described with some alterations (25, 26). Rundown microsomes were pre-incubated in binding reaction mixture lacking NTPs for 15 minutes at 32°C before being added to binding reactions containing NTPs, rat liver cytosol and the indicated concentrations of Sar1 proteins(26). Control microsomes were not pre-incubated in any way. All reactions were incubated at 32°C for 15 min and terminated by transfer to ice for 10 min. Membranes were salt washed and collected by centrifugation at 16,000 g for 15 min at 4°C, and membrane pellets were resuspended and resolved on SDS-PAGE gels for Western blot analysis(17, 25, 31). The same blots were probed with anti Sec23 and Sec31 -specific antibodies. All presented assays were repeated at least 3 times with the exception of the PI4P addition (Fig. 2D) which was done twice.

### Rundown Morphological Recruitment

The morphological analysis of coat assembly and tubulation of ERES in semi-intact HeLa cells were performed as previously described with some alterations (17, 25, 42). In brief, HeLa cells were permeabilized with 40 µg/ml digitonin in KHM buffer for 5 min on ice, washed with KHM, then rundown reactions were pre-incubated in reaction mixture lacking NTPs for 15 minutes at 32°C before incubation with transport cocktail (26) in the presence of rat liver cytosol at 32°C for 30 minutes to measure coat assembly. Control reactions were not pre-incubated following KHM washes but were directly incubated with transport cocktail. All cells were fixed in 4% formaldehyde in PBS for 15 min at RT for coat assembly analysis with Sec13, Sec24c, Sec31, and Sec31a specific antibodies. All morphological assays were repeated at least 3 times with similar outcomes. Permeabilized cells were labeled and processed for EM as previously described (3, 41). Indirect immunofluorescence was carried out as described previously (26).

### Imaging and analysis

Images (for fixed or live imaging) were acquired on an Olympus Fluoview 1000 confocal system using an inverted microscope (IX-81 Olympus) and 60×NA 1.42 PLAPON objective. Images were collected using FV10-ASW V. 02.00.03.10 (Olympus Corporation), sized and incorporated in figures without further processing using Adobe Illustrator CC and Canvas X software. STED microscopy was conducted on a Leica SP8 STED 3× microscope (Leica Microsystems, Wetzlar, Germany) equipped with a SuperK EXTREME EXW-12 pulsed white light laser (NKT Photonics Inc., Portland, OR) as an excitation source, a Katana-08HP pulsed depletion laser as a depletion light source (775 nm; NKT Photonics), and a Leica high-contrast plan apochromat 93× 1.30 numerical aperture (NA) glycerol CS2 objective with motorized correction collar (Leica Microsystems); imaging was controlled by Leica Application Suite X software (LAS X, Leica Microsystems). Immuno gold staining and electron microscopy was conducted as previously described(3, 41). For quantitative analysis, experiments (comprised of randomly collected confocal images of cell fields) were subjected to a constant threshold per experiment. Experiments were analyzed for overall intensities of collagen 1 in cells using Image J (Fiji). Similar collection of confocal images of cell fields with constant threshold application was utilized to define Manders’ Overlap Coefficient using the Fiji plugin JACoP. All data were subjected to statistical analysis using two tailed T-test.

### Membrane Contact Assay

HeLa microsomes for rundown reactions were pre-incubated in binding reaction mixture for 15 minutes at 32°C. Control HeLa microsomes were not. Membranes were pelleted at 16,000g for 3 minutes at 4°C [MSP]. Pellets and an aliquot of supernatant were saved for analysis. Half of each supernatant was pelleted at 54,000rpm (179000g) for 20 minutes at 4°C in an ultracentrifuge [HSP]. All pellets and supernatants were resolved on 12% SDS-PAGE gels for Western blot analysis with ERGIC-53 or Sec12 specific antibodies. Western blots were quantified by densitometry using Image J. The experiment was repeated 3 times with similar results.

### GST-Pulldowns

250µg of GST, GST-HsSec31a1041-1220, or GST-HsSec31a1041-1113 were incubated with Glutathione Sepharose^TM^ 4B (GE Healthcare) beads at 4°C for 1-2h. Unbound GST protein was removed by washing 3 times with 1X Assay Buffer [25mM HEPES, pH7.2; 125mM KOAc; 5mM MgOAc; 2mM EDTA; 2mM EGTA]. HeLa cells expressing pEGFP-p125A, were lysed with 1X Assay Buffer + 1% TritonX-100. Two thirds of a 100mM dish were used for each pulldown reaction. The GST-protein-bound beads were incubated with HeLa lysates at 4°C for 1 hour. Beads were washed 2 times with 1X Assay Buffer and 2 times with 1XPBS. The beads were resuspended in 2X sample buffer and the proteins resolved on SDS-PAGE gels for Western blot analysis with p125A-specific antibodies. PD assays were repeated 3 times with similar results.

### Gene-Editing in U2OS cells

The SEC23IP (p125A) knock out U2OS cell line was created using Crispr-Cas9 genome editing using one gRNA at the 5’ end of the SEC23IP gene (sg1: gaagtaactaaatgtctgtg) and one at the 3’ end of the gene (sg2: ggagaacgtagtcaattcgg) and a single stranded oligo guiding repair of the site favoring deletion of most of the coding region of the gene (ssODN: aggacagcttccttggtcagacttctattcacacatctgtgacacaattgactacgttctccaagaaaaaccaatagagagttt taatga). SEC23IP gRNAs were cloned into the BbsI site of px458 (addgene 48138) or px458-mCherry (the latter having the EGFP gene of px458 replaced by mCherry) after annealing of the oligo’s (sg1: caccgGAAGTAACTAAATGTCTGTG and aaacCACAGACATTTAGTTACTTCc cloned into px458-mCherry; sg2: caccgGGAGAACGTAGTCAATTCGG and aaacCCGAATTGACTACGTTCTCCc cloned into px458). For transfection U2OS cells were trypsinized, counted and transfected while seeding with 500 ng or each gRNA expressing plasmid and 400 pmol of ssODN. Three days after transfection transfected cells were FACS sorted as single cells into 96 well plates for further screening. Single cell clones were expanded and screened by PCR for presence of the deletion fragment and absence of the wild type fragments using the following primers: deletion fragment ctgcggagtgatagagtaccc and tgctagcaccaaaagcccac; wild type fragment around sg1 ctgcggagtgatagagtaccc and gagggagactacttggctgag; wild type fragment around sg2 gcctggtagctcctcaggttg and tgctagcaccaaaagcccac. To confirm the presence of only the deleted gene in the selected clone we performed transcript analysis by amplifying SEC23IP (p125A) transcripts from cDNA using primers upstream and downstream of the gRNAs therefore amplifying both wild-type and deleted transcripts. RNA was isolated using the NZY Total RNA Isolation kit (NZYTech) and cDNA was prepared using the SuperScript III Reverse Transcriptase (Invitrogen). Transcripts were amplified using primers ccatggccgagagaaaacctaac and tcaatgctggggctgttctgg. Sequencing analysis of the obtained transcripts further confirmed the deletion of the gene in 2 different deletion alleles. Absence of the protein in the selected line was established by western blotting using a SEC23IP antibodies (HPA038403,).

### Sec31 stability in control and KO cells

For the analysis of Sec31A stability in parental or p125A deleted U2OS cells, cells were washed with ice cold PBS and permeabilized on ice using KHM buffer supplemented with 0.1% saponin (W/V). Cells were washed once with KHM fixed and stained with indicated antibodies.

### Collagen traffic assays

Collagen accumulation of and traffic analysis was conducted as previously reported (72). Briefly cells were washed 3 times with PBS and incubated in DMEM 1%FCS for 3 hrs at 40°C under standard incubation environment (humidified, 5% CO2). Cells were supplemented with 100μg/ml cycloheximide, 50μM ascorbic acid and 25mM Hepes-KOH (pH 7.2) and incubated for defined periods at 32°C, before transfer to ice, washes (3X ice cold PBS) and fixation for imaging. Cells were fixed and stained for immunofluorescence analysis.

### tsVSV-G traffic assays

Parental and U2OS,Δp125A cells were transfected with a plasmid encoding tsVSV-G-GFP using X-tremeGENE-R according to manufacturer’s protocol with optimized DNA concentration. The cells were incubated for 5 hr. at 37°C, DMEM, 10% FCS media was added and the cells were moved to an incubator set at 40°C for an overnight incubation (18 hrs). Media was replaced with Optimem supplemented with 100μg/ml cycloheximide and 25mM HEPES pH-7.4 and cells were incubated at 32°C for indicated times. Cells were fixed and processed for indirect immunofluorescence using indicated antibodies directed at Golgi markers and GFP. For live imaging, tsVSV-G transfections were carried by electroporation using the Neon electroporation system (Invitrogen) according to manufecturer’s protocol optimized for U2OS cells. Transfected cells were immediately plated on slides (Mattek corp), incubated at 37°C, for 40 hrs, and transferred to 40°C for an overnight incubation in DMEM, 10%FCS. The cells were washed with ice cold PBS, incubated on a temperature-controlled microscope stage at 32°C in Optimem, 100μg/ml cycloheximide and 25mM HEPES pH-7.4 and imaged continually for indicated times.

### Antibodies

Polyclonal and monoclonal antibodies against Sec31A, polyclonal anti Sec13, Sec12, Sec23, Sar1 and Sec24C antibodies were kindly provided by Dr. Bill Balch (TSRI, La Jolla CA). Antibody to Sec24C from Invitrogen (#PA5-59101), antibody to Sec31A was from BD Transduction Laboratories (#612351). COPII subunit antibodies were re-verified by functional recruitment assays (using indirect immunofluorescence) and/or by using western blots to follow expression or depletion of target antigens. Antibodies against p125A were from Bethyl Labs (#MSTP053) and Atlas antibodies (#HPA038403). Antibody against collagen 1 from Abcam (#ab138492). Anti ERGIC53 was from Anzo Life Sciences (ALX-804-602), Antibodies to Giantin and GPP130 were kindly provided by Dr. A. Linstedt (Carnegie Mellon U. Pittsburgh PA), anti-GRASP65 (C-20) from Santa Cruz (# sc-1948). Anti FLAG antibodies (M2, F1804 Sigma), antibodies to Sec16A, Novus Biologicals (NB100-1799), antibodies to p97/VCP, RDI research diagnostics (RDI PR065278), antibodies to GFP, Polysciences Inc. (24240) and Proteintech (66002-1-ig). Anti-human CD25(IL2-R alpha, TAC) from Ancell immunology research products (174–200).

### Proteins

Recombinant Sar1-H79G protein was produced and purified as previously described (26, 73). GST and GST tagged hSec31A protein fragments were purified on GS-Sepharose 4B (GE Healthcare Life Science).

### Omics

Approximately, one million Control U2OS and p125A KO U2OS were seeded in each 60mm tissue culture dish with DMEM, high glucose, GlutaMAX™ Supplement, pyruvate (Gibco; 31966021) containing 10% foetal bovine serum (Gibco; A5256701), 1% penicillin-Streptomycin (Gibco;15140122) and 0.25 mM 2-Phospho-L-ascorbic acid trisodium salt (Sigma; 49752). A day before the experiments, cells were washed three times in PBS and incubated in 5ml of the same medium without serum for 18 hours.

### Transcriptomics

total RNA isolation was performed using the NucleoSpin RNA Kit (Macherey-Nagel) according to manufacturer’s recommendations. Approximately 50 μL RNA eluates were obtained. Concentration and RNA integrity number (RIN) were assessed using DeNovix and RNA ScreenTape assay (Agilent, 5067-5576). RNA library preparation and sequencing were performed by Novogene Co. Ltd. (United Kingdom) using human strand-specific mRNA (WBI-Quantification), directional mRNA library preparation (poly A enrichment) and 2 × 150 bp pair-end configuration (PE) with raw data of 8.0 Gb per sample. The prepared libraries were subsequently pooled for sequencing on the NovaSeq platform (Illumina, San Diego, CA, USA), according to optimal library concentration and data amount. All samples were sequenced in the same batch. During the data quality control process, raw reads of the FASTQ format were processed before clean reads were obtained for downstream analyzes. Novogene conducted quantification of gene expression levels, co-expression analysis, and enrichment analysis based on whole transcriptome RNA-Seq data.

### Proteomics

after adding protease and phosphatase inhibitors, medium was subjected to centrifugation at 2500g at 4°C for 20 minutes to remove any cells or cellular debris. Medium was then concentrated using centrifugal filter units (Millipore; UFC801024) with a 10 kDa cut off. Approximately, 50µl of concentrated medium was frozen and stored at - 80°C till further use. Intracellular proteins were collected using RIPA buffer (SERVA; 39244) according to the manufacturer’s instructions. Protein concentrations were determined using a Pierce™ BCA Protein Assay Kit (Thermo Scientific; 23225).

Mass spectrometry grade chemicals were used, including acetonitrile (ACN), H2O, ammonium bicarbonate (NH4HCO3), formic acid (FA), Tris(2-Carboxyethyl) Phosphine Hydrochloride (TCEP-HCl) and S-Methyl methanethiosulphonate (MMTS) (ThermoFisher Scientific). Sequencing grade Trypsin/Lys C mix was from Promega. Ammonium bicarbonate (NH4HCO3) was from Sigma-Aldrich. For sample preparation, a six-fold volume of cold acetone (−20°C) was added to each sample volume containing 10 µg of protein extracts. Vortexed tubes were incubated overnight at −20°C then centrifuged for 10 min at 11,000 rpm at 4°C. Supernatant was removed, then the protein pellets were dissolved in 8M urea, 25 mM NH4HCO3 buffer. Samples were then reduced with 10 mM TCEP-HCl and alkylated with 20 mM MMTS. After a 16-fold dilution in NH4HCO3, samples were digested overnight at 37°C by a mixture of trypsin/Lys C (1/20 Enzyme/Substrate ratio). The digested peptides were loaded and desalted on Evotips Pure, provided by Evosep one (Odense, Denmark) according to manufacturer’s procedure.

For LC-MS/MS acquisition, samples were analyzed on a timsTOF Pro 2 mass spectrometer (Bruker Daltonics, Bremen, Germany) coupled to an Evosep one system (Evosep, Odense, Denmark) operating with the 30SPD method developed by the manufacturer. Briefly, the method is based on a 44-min gradient and a total cycle time of 48 min with a C18 analytical column (0.15 x 150 mm, 1.9µm beads, ref EV-1106) equilibrated at 40°C and operated at a flow rate of 500 nL/min. H2O/0.1 % FA was used as solvent A and ACN/ 0.1 % FA as solvent B. The timsTOF Pro 2 was operated with a DIA-PASEF method comprising 12 pydiAID frames with 3 mass windows per frame resulting in a cycle time of 0.975 seconds as described in Bruker application note LCMS 218.

Raw MS data files were processed using Spectronaut 18 (Biognosys, Switzerland). Data were searched against the SwissProt Mus Musculus (06 2024, 17219 entries) + Bovine (06 2024, 6046 entries) databases. Contaminating proteins identified only in the Bovine taxonomy were excluded from the final protein list. Specific tryptic cleavages were selected and a maximum of 2 missed cleavages were allowed. The following post-translational modifications were considered for identification: Acetyl (Protein N-term), Oxidation (M), Deamidation (NQ) as variable and MMTS (C) as fixed. The maximum number of variable modifications was set to 3. Identifications were filtered based on a 1% precursor and protein Q-value cutoff threshold. The protein LFQ method was set to automatic and the quantity was set at the MS2 level with a cross-run normalization applied. Multivariate statistics on protein measurements were performed using Qlucore Omics Explorer 3.9 (Qlucore AB, Lund, Sweden). A positive threshold value of 1 was set to allow a log2 transformation of abundance data for normalization, i.e. all abundance data values below the threshold are replaced by 1 before transformation. The transformed data were finally used for statistical analysis, i.e. the evaluation of differentially present proteins between two groups using a two-sided Student’s t-test. An adjusted p<0.05 was used to filter differential candidates.

## Author Contributions

KRL: conceptualization, experimental work, analysis and editing GS: Experimental work data quantifications and analysis NB: experimental work and analysis, MV: experimental work and analysis, IR, VM: analysis writing and editing. MA: conceptualization, supervision, experimental work, analysis, writing and editing.

## Acknowledgments

We thank Drs W. E. Balch, R. Pepperkok, A. Linstedt, D. C. Devor, L. M. Traub and J. Lippincott-Schwartz for generously providing valuable reagents for the study. We thank the CRG advanced light microscopy unit (ALMU). We thank the Center for Biological Imaging (CBI) at the University of Pittsburgh School of Medicine for help with transmission EM and STED microscopy. We thank the proteomics facility (ProteoSeine) of the Institut Jacques Monod for assistance with the mass spectrometry. The study was initially supported by NIH grant 5R01DK092807 (MA). We further acknowledge the support of the Spanish Ministry of Science and Innovation to the EMBL partnership, the Centro de Excelencia Severo Ochoa (CEX2020-001049, MCIN/AEI/10.13039/501100011033) the Generalitat de Catalunya through and the CERCA Programme and to the EMBL partnership. We are grateful to the CRG Core Technologies Programme for their support and assistance in this work. We acknowledge financial support from the following sources: Ministerio de Economía y Competitividad (PID2019-105518GB-100 to VM) and MatSec, from Agence Nationale de la Recherche and AJE202210016216 from Fondation Recherche Medicale (IR).This project has received funding from the European Research Council (ERC) through the European Union’s Horizon 2020 research and innovation programme under grant agreement 951146-LiquOrg.

---

**Supplement Figure 1.**
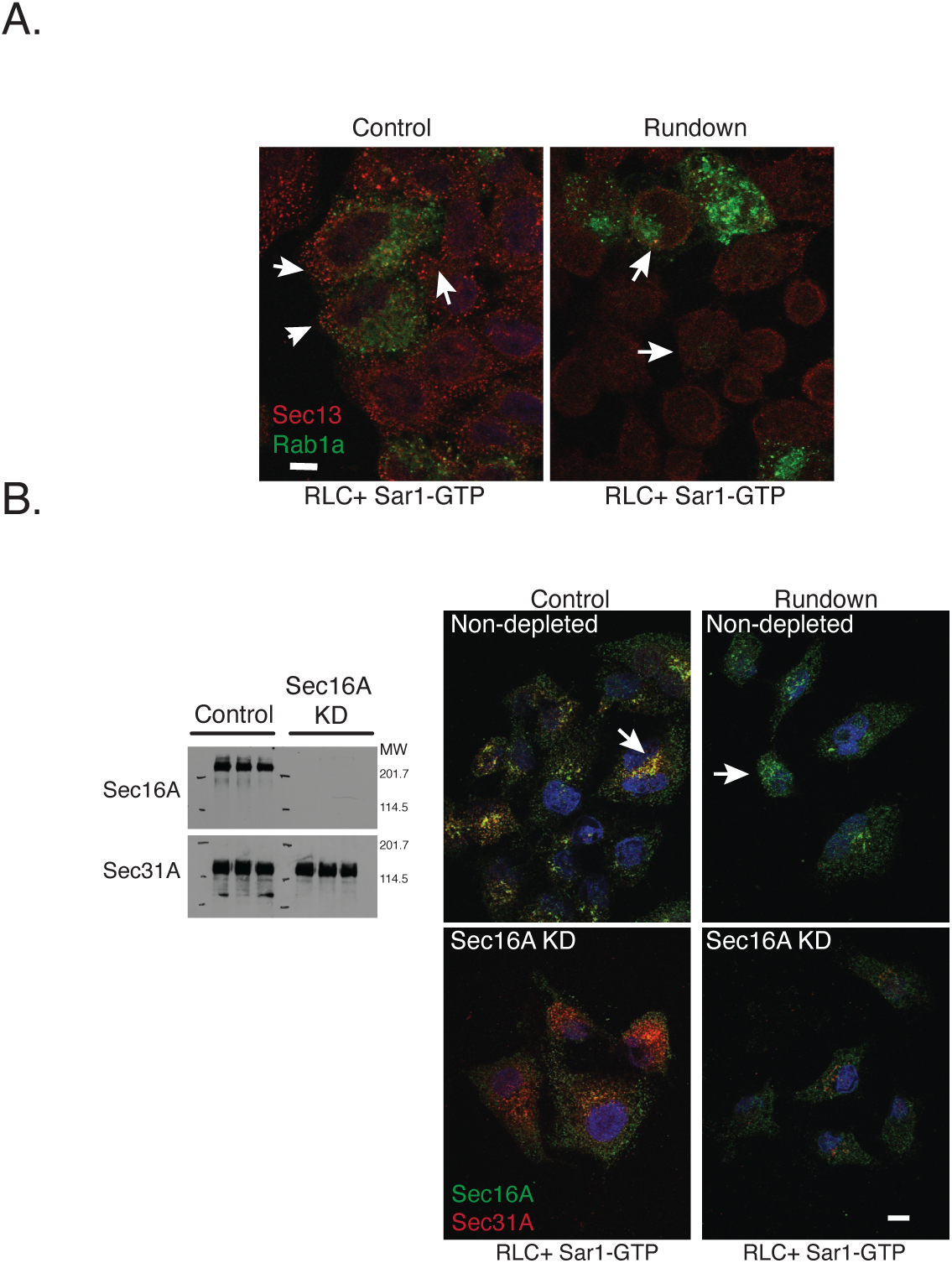
Active Rab1 or Sec16A depletion do not rescue the linkage between COPII inner and outer layers. **A.** Cells were transfected with construct expressing Flag-Rab1a-GTP(Q70L), permeabilized and subjected to rundown and COPII recruitment as in Fig. 1, monitoring Sec13 (red) and Rab1-GTP (anti Flag, green). Note that Rab1a-GTP is retained in cells and occasionally preserves Sec13 at the sites. **B.** Sec16A was depleted using siRNA and analyzed by western blot with Sec31A serving as a control as indicated (triplicates of control or KD samples are shown in the left panel). Depleted cells were permeabilized and subjected to rundown pre-incubation. Subsequently, control or Sec16A depleted cells were supplemented with RLC and SAR1-GTP (as in Fig.1). The recruitment of the coat outer layer (Sec31A, monoclonal antibody) and Sec16A localization was determined by IF. Bars are 5 μm.

**Supplement Figure 2.**
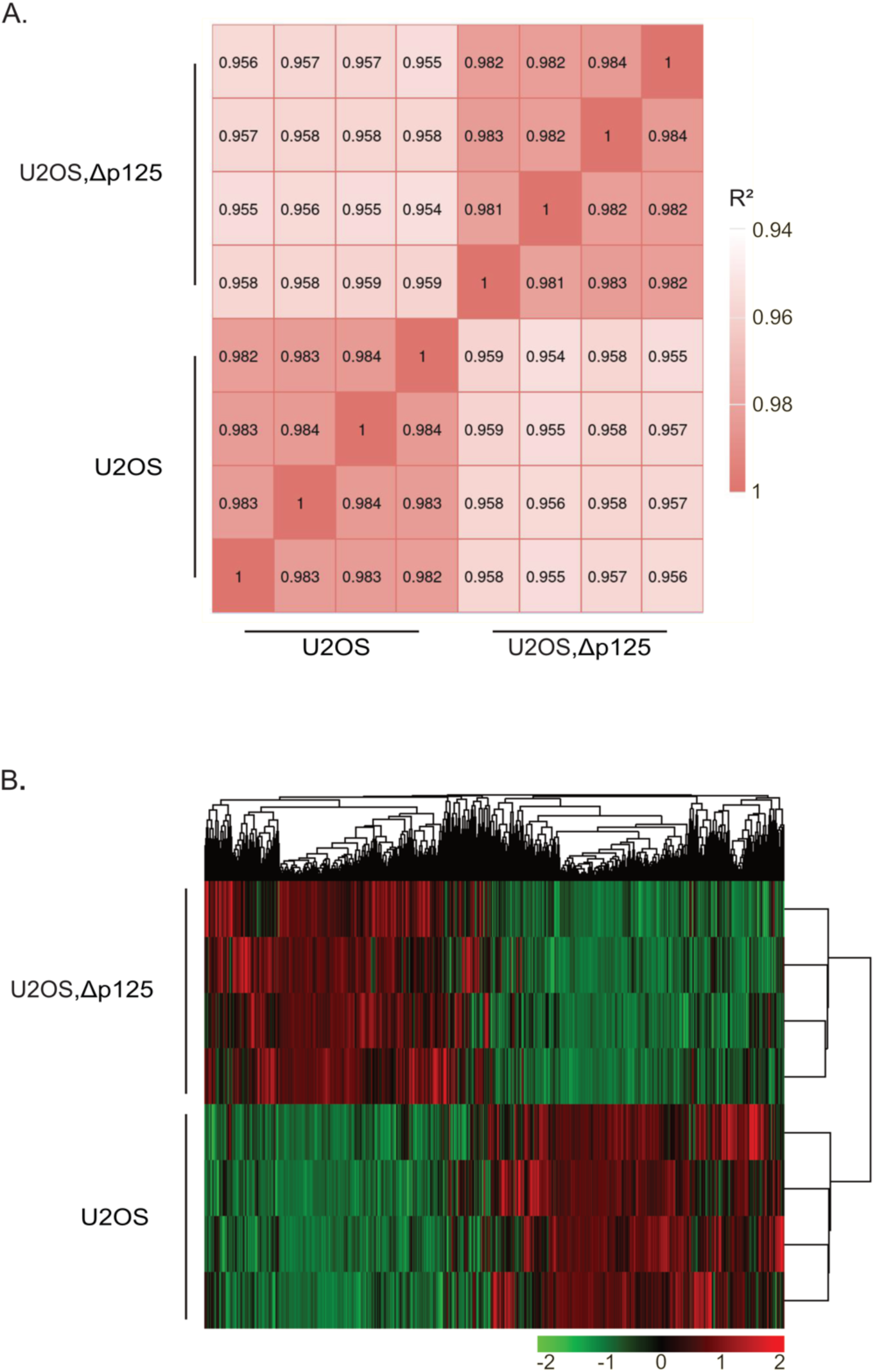
**A.** Heatmap of Pearson correlation coefficient matrix. Correlation coefficient between the samples and conditions plotted as heat maps. The higher the Pearson correlation coefficient (R²) of the sample is, the closer the expression pattern. The bar on the right of the figure indicates the color legend of R² calculated for each pair of samples in the matrix. The closer the correlation coefficient is to 1, the higher similarity between samples. Encode suggests that R² should be greater than 0.92 (ENCODE Project Consortium, 2004). **B**. Heatmap of differentially expressed genes (padj < 0.05 and |log 2 (FoldChange)| > 1) identified between the U2OSΔp125 and U2OS. Genes or samples with similar expression patterns in the heat map cluster together. The color in each grid reflects not the gene expression value, but the value obtained after homogenizing the expression data rows (generally between - 2 and 2). Therefore, the colors in the heat map can only be compared vertically (the expression of the same gene in different samples), but not horizontally (the same sample). There are not only inter-group clustering, but also inter-sample clustering.

**Supplement Figure 3.**
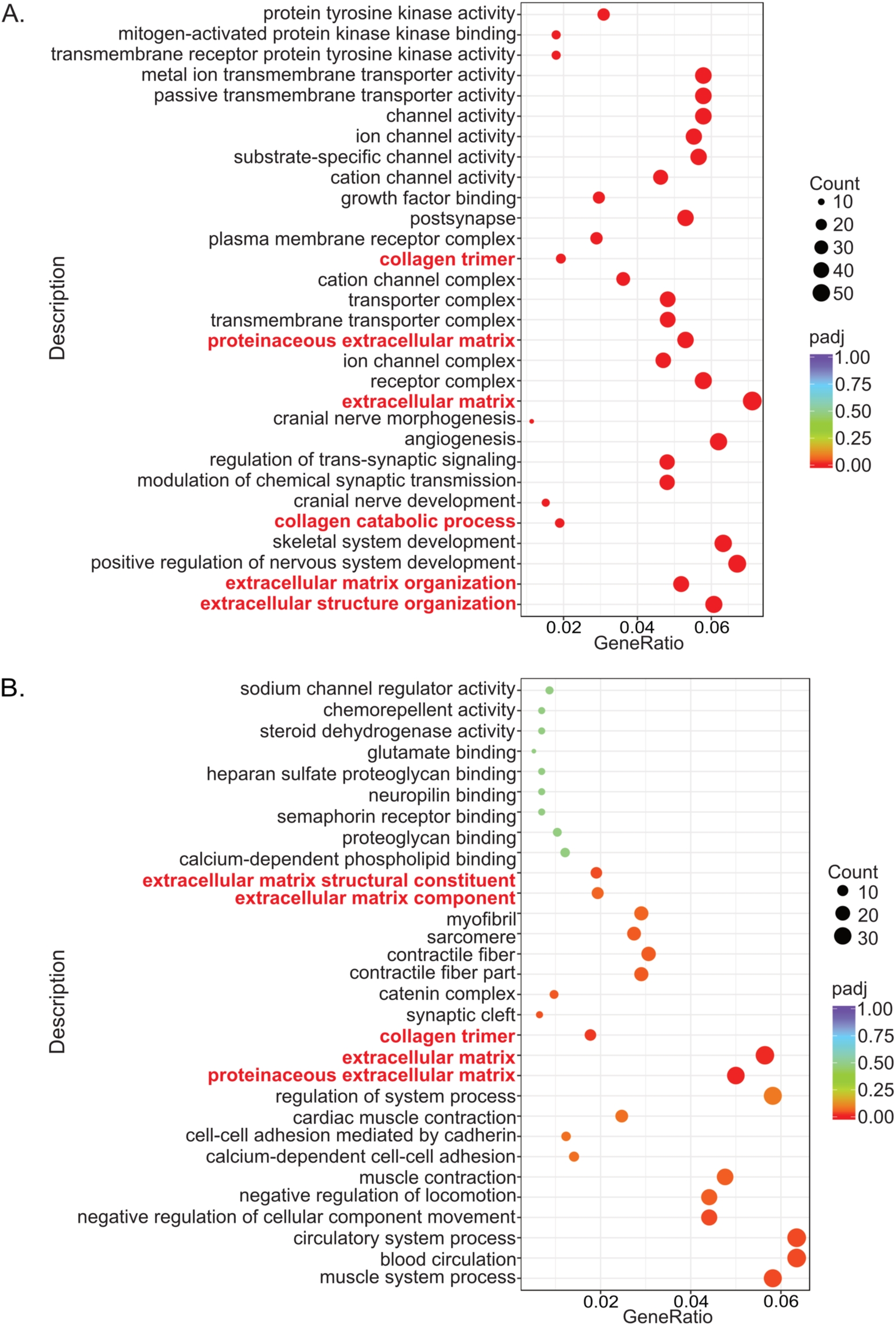
Dot plot of the Gene Ontology (GO) enrichment analysis for down regulated genes (**A.**) and up regulated genes (**B.**). The most significant 30 GO Terms were selected for display. The x-axis is the ratio of the number of differential genes linked with the GO Term to the total number of differential genes, and the y-axis is GO Term. The size of a point represents the number of genes annotated to a specific GO Term, and the color from red to purple represents the significant level of the enrichment. GO pathways with padj < 0.05 are significant enrichment. (See Supplementary Table 1 for complete genes lists).

**Supplement Figure 4.**
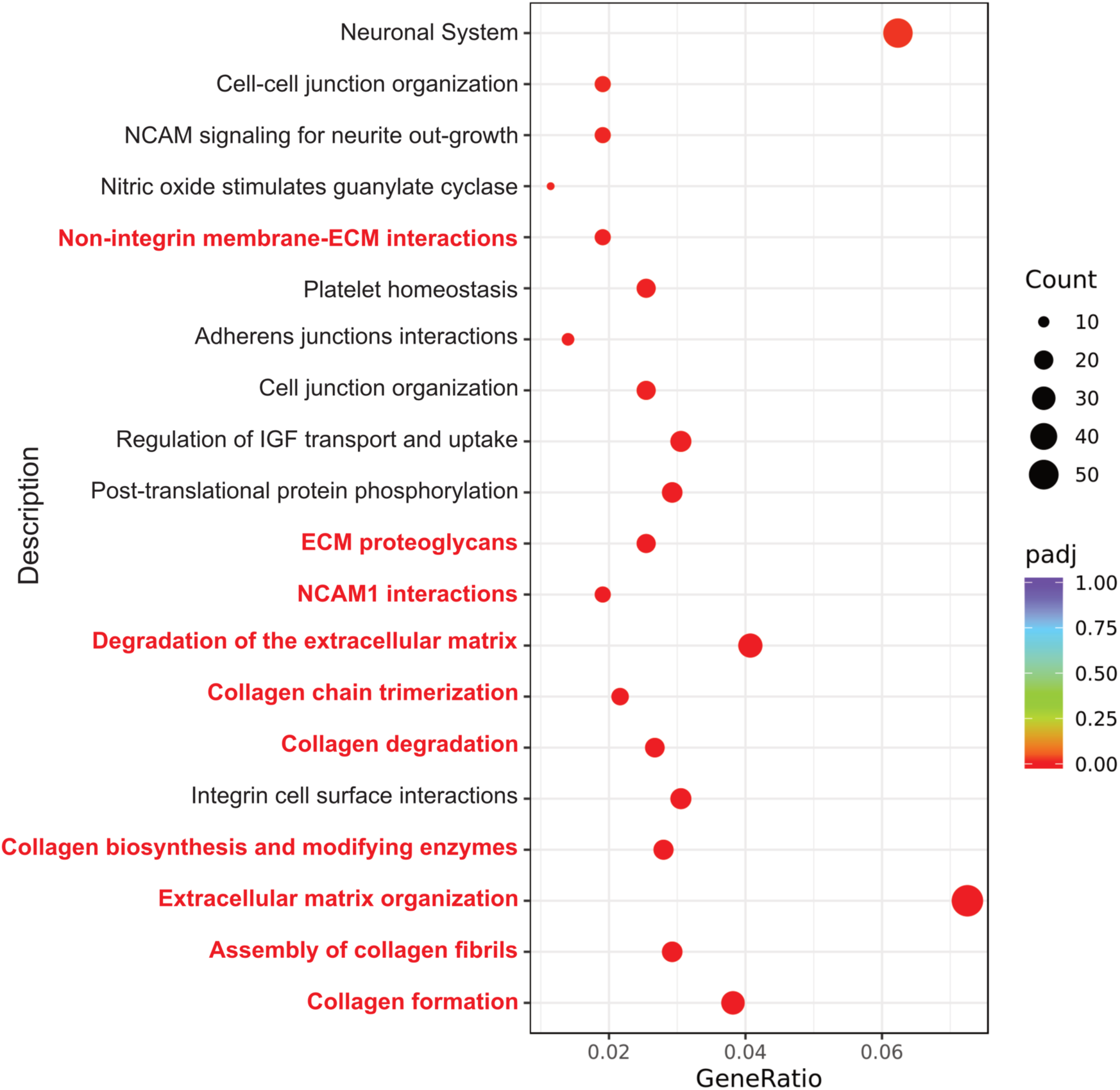
Reactome enrichment analysis. The most significant 20 Reactome pathways were selected for display. The x-axis is the ratio of the number of differential genes linked with the Reactome pathway to the total number of differential genes, and the y-axis is Reactome Pathways as listed. The size of a point represents the number of genes annotated to a specific Reactome pathway, and the color from red to purple represents the significant size of the enrichment. (See Supplementary Table 1 for complete genes lists).

**Supplement table1. Transcriptomics Data (Excel attached).**

1. The full list of differentially expressed genes identified between U2OSΔp125 and U2OS. 2. List of significantly up regulated expressed genes identified in U2OSΔp125 compared to U2OS. 3. List of significantly down regulated expressed genes identified in U2OSΔp125 compared to U2OS group.4. List of differentially expressed genes involved in the most significant 30 GO Terms. 5. List of down regulated expressed genes in U2OSΔp125 involved in the most significant 30 GO Terms. 6. List of up regulated expressed genes in the U2OSΔp125 involved in the most significant 30 GO Terms. 7. List of differentially expressed genes involved in the most significant 20 Reactome pathways.

**Supplement Table 2.**
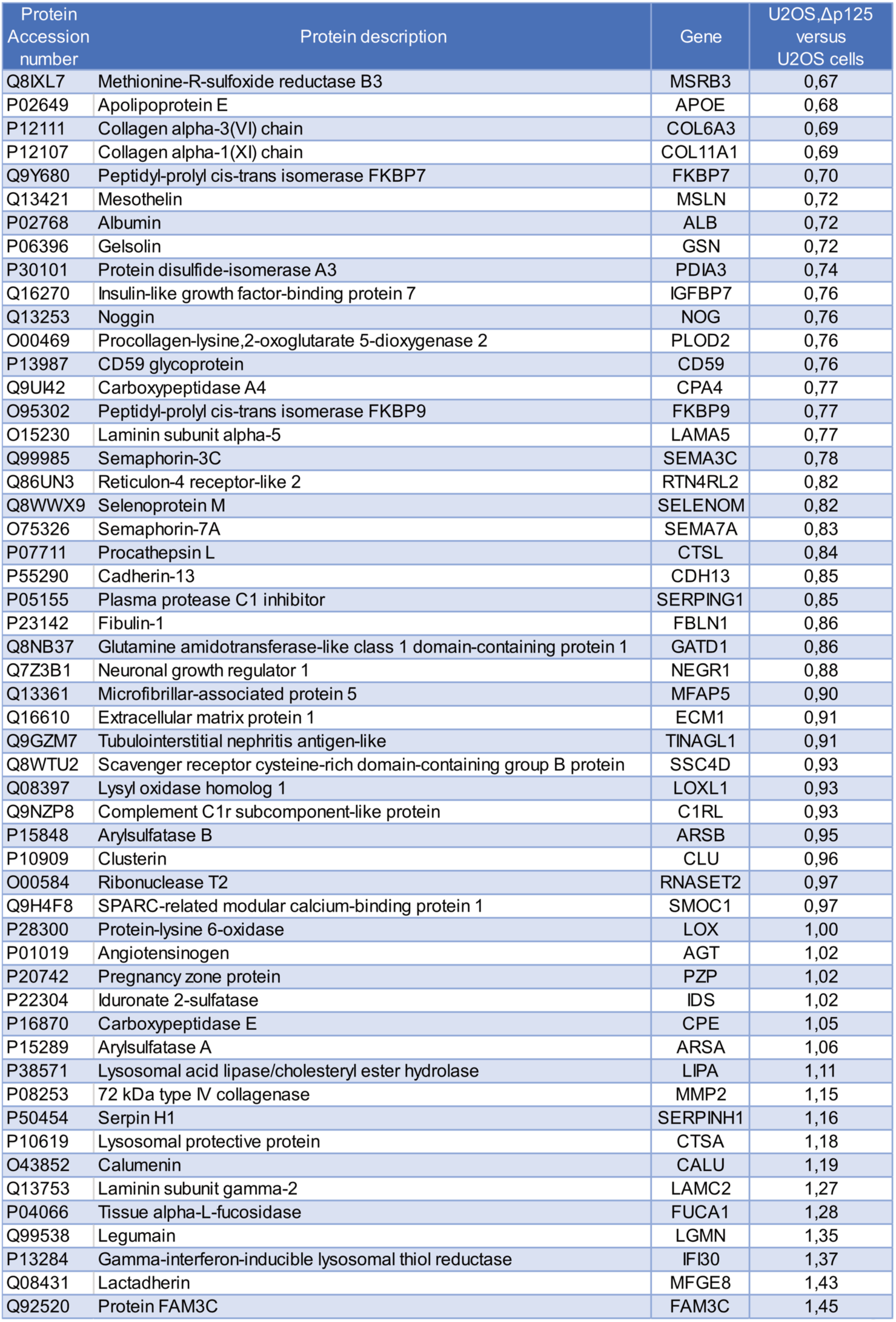
List of proteins with unchanged secretion in U2OSΔp125 compared to U2OS. Secreted and intracellular proteins were quantified by LC-MS/MS in U2OSΔp125 and U2OS. Within each group, the proportion of the secreted protein detected was determined. Then, each value for U2OSΔp125 was compared to U2OS. 54 proteins were identified with unchanged secretion (threshold between 0.67 and 1.50). (See Supplementary Table 4 for complete proteins lists detected in lysate and secretome).

**Supplement Table 3.**
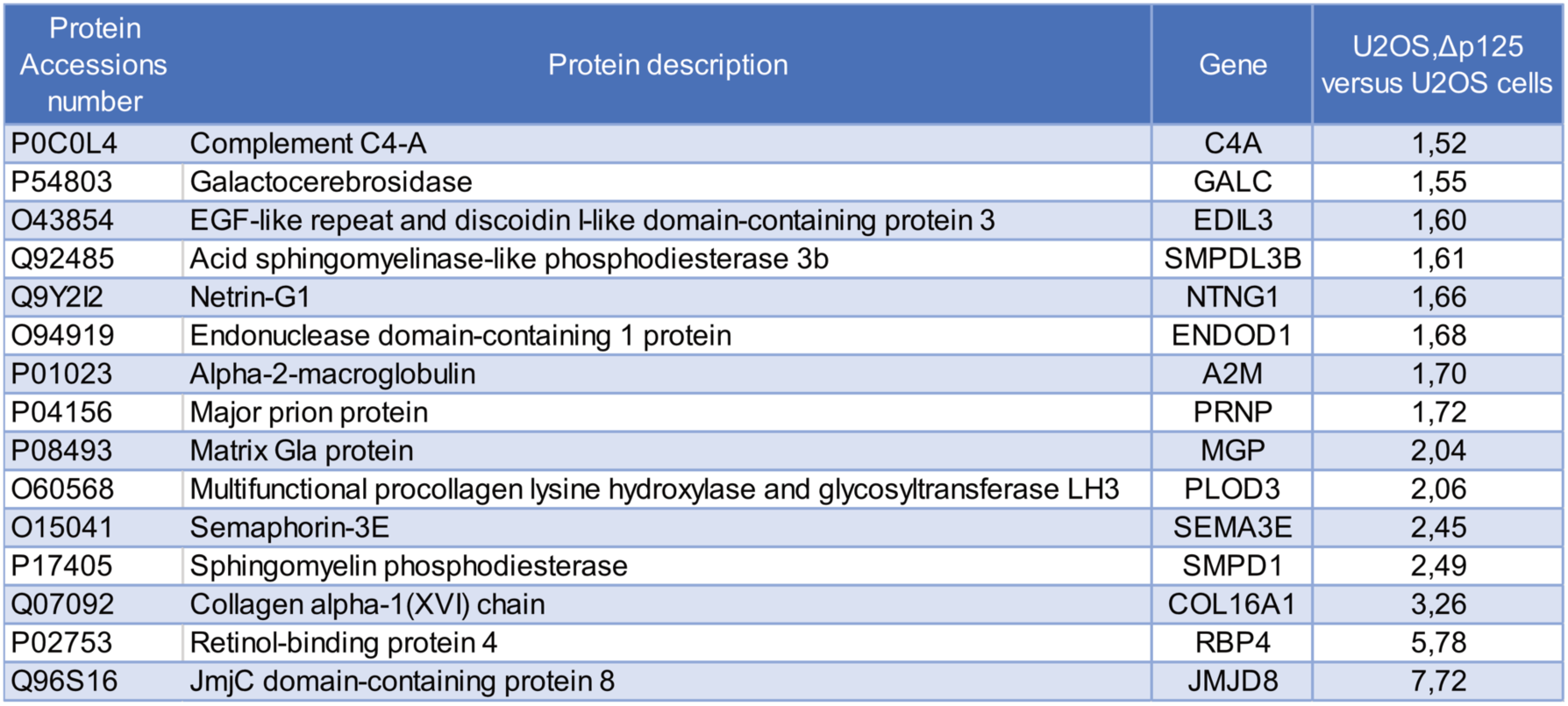
List of proteins identified as hypersecreted in U2OSΔp125 compared to U2OS. Secreted and intracellular proteins were quantified by LC-MS/MS in U2OSΔp125 and U2OS. Within each group, the proportion of the secreted protein detected was determined. Then, each value for U2OSΔp125 was compared to U2OS. 15 proteins were finally identified as hypersecreted with a threshold set at 1.50. (See Supplementary Table 4 for complete proteins lists detected in lysate and secretome).

**Supplement Table 4.** (Excel attached).

Proteomics data. 1, The complete list of intracellular proteins identified and quantified in U2OSΔp125 and U2OS. Fold changes were calculated. P values were calculated using Student’s t test. 2. The complete list of secreted proteins identified and quantified in U2OSΔp125 and U2OS. Fold changes were calculated. P values were calculated using Student’s t test.

**Supplement Movie 1.** U2OS cells expressing tsVSV-G-GFP were transferred from 40°C to 32°C with added cycloheximide (materials and methods) and directly imaged at mid cell at a rate of 7.05sec/frame, recording time is 20 min.

**Supplement Movie 2.** U2OSΔp125A cells expressing tsVSV-G-GFP were transferred from 40°C to 32°C with added cycloheximide (materials and methods) and directly imaged at mid cell at a rate of 7.05sec/frame, recording time is 20 min.

